# SOURSOP: A Python package for the analysis of simulations of intrinsically disordered proteins

**DOI:** 10.1101/2023.02.16.528879

**Authors:** Jared M. Lalmansingh, Alex T. Keeley, Kiersten M. Ruff, Rohit V. Pappu, Alex S. Holehouse

## Abstract

Conformational heterogeneity is a defining hallmark of intrinsically disordered proteins and protein regions (IDRs). The functions of IDRs and the emergent cellular phenotypes they control are associated with sequence-specific conformational ensembles. Simulations of conformational ensembles that are based on atomistic and coarse-grained models are routinely used to uncover the sequence-specific interactions that may contribute to IDR functions. These simulations are performed either independently or in conjunction with data from experiments. Functionally relevant features of IDRs can span a range of length scales. Extracting these features requires analysis routines that quantify a range of properties. Here, we describe a new analysis suite SOURSOP, an object-oriented and open-source toolkit designed for the analysis of simulated conformational ensembles of IDRs. SOURSOP implements several analysis routines motivated by principles in polymer physics, offering a unique collection of simple-to-use functions to characterize IDR ensembles. As an extendable framework, SOURSOP supports the development and implementation of new analysis routines that can be easily packaged and shared.

## 1. INTRODUCTION

Natively unfolded proteins or intrinsically disordered proteins and regions (collectively referred to as IDRs,) are a ubiquitous class of proteins and domains that regulate a variety of molecular functions and cellular phenotypes^1–4^. Unlike folded domains, which are well-described by a small number of structurally similar microstates, IDRs are defined by their conformational heterogeneity ^4,5^. As a result, the accurate description of IDRs in the solution states necessitates a statistical description of the underlying conformational ensembles^6^. These ensembles, which are affected by changes to solution conditions and the types of components present in the solvent, are distributions of energetically accessible protein configurations that capture the sequence-encoded conformational biases associated with a given IDR ^4,7,8^. Several studies have established direct connections between sequence-ensemble relationships of IDRs and the molecular functions of these conformationally heterogeneous regions ^8–10^. Accordingly, there is a need for facile, ready-to-use methods to uncover the molecular grammar that underlies sequence-ensemble-function relationships of IDRs. ^9^.

Measurements of IDR ensembles in solution allow for quantitative mapping of sequence-ensemble relationships. Techniques that obtain statistical information on molecular configuration without assuming a single dominant state are well-equipped to characterize IDR ensembles. These techniques include static and dynamic light scattering (SLS and DLS, respectively), small-angle X-ray scattering (SAXS), circular dichroism (CD), nuclear magnetic resonance (NMR) spectroscopy, multiparameter fluorescence spectroscopies, and other single-molecule techniques ^7,11–16^. While these experimental techniques offer a window into conformational behavior, they typically probe a single class of molecular configuration (*e.g*., global ensemble average dimensions, distances between specific positions along the chain, *etc*.). Alongside these experimental approaches, all-atom and coarse-grained molecular simulations are routinely deployed to make predictions or interpret data obtained from experimental measurements. This coupling of experimental and computational methods allows for the integration of multiple conformational inputs, enabling a holistic assessment of sequence-ensemble mapping ^17–23^.

Simulations of all stripes, but specifically all-atom simulations based on explicit or implicit representations of solvent are especially useful for describing sequence-specific conformational ensembles of IDRs^24,25^. If a simulation can fully explore the conformational landscape and the forcefield being used is accurate enough, then all-atom molecular simulations enable the direct prediction of ensembles from sequence. These computationally derived ensembles can be compared directly or indirectly with experiments or used in isolation to understand functional and evolutionary constraints on IDRs ^19,20,25^. Consequently, there has been substantial interest in developing and applying Molecular Dynamics (MD) and Monte Carlo (MC) simulations to study IDRs ^26–33^.

As all-atom simulations have become increasingly routine, various software packages have emerged to perform and analyze molecular simulations. Major packages for performing all-atom simulations (so-called simulation engines) include, but are not limited to, Amber, CAMPARI, CHARMM, Desmond, GROMACS, LAMMPS, OpenMM, and NAMD ^30,34–40^. Alongside the development of simulation engines, there has also been an emergence of stand-alone packages for simulation analysis. Although most simulation engines contain their analysis routines, stand-alone analysis packages provide an alternative that, in principle, can be relatively lightweight, customizable, and unburdened by coding practices or conventions of the inevitably larger simulation engines. General-purpose analysis packages include Bio3D, CPPTRAJ, ENSPARA, LOOS, MDAnalysis, MDTraj, ST-Analyzer, VMD, and others^41–43^ (see **Supplemental Table 1** for a more extensive list). While some packages are general-purpose libraries for analyzing simulation trajectories, others are developed with a specific goal in mind ^44–46^. The ability to decouple analysis from performing simulations allows for ease of use, installation, and portability to be prioritized in analysis packages, while performance can be prioritized in simulation engines. It also enables familiarity with a single analysis framework that can be applied across different simulation engines.

All-atom simulations of IDRs are becoming increasingly common ^17,19,47^. Despite this, there is a lack of stand-alone analysis packages that specifically cater to the analysis of IDR conformational ensembles. Given their inherently heterogeneous ensembles and the lack of a relevant single reference structure, many of the structure-centric analyses commonly employed in the context of folded may be poorly-suited for characterizing IDR ensembles. In contrast, concepts and principles from polymer physics have been taken and applied to interpret and understand disordered and unfolded proteins to great effect ^6,11,19,48–51^.

Here we introduce SOURSOP (**S**imulation analysis **O**f **U**nfolded **R**egion**S O**f **P**roteins), a Python-based software package for the analysis of all-atom simulations of disordered and unfolded proteins. SOURSOP combines both analysis routines commonly found for folded proteins with a range of IDR-centric analyses that have been used to great effect across many publications over the last half-decade. In the remainder of this article, we lay out the software architecture of SOURSOP, provide several examples of analysis that can be performed, and offer a discussion of practical and conceptual features associated with the software.

## 2. METHODS

SOURSOP is written in Python 3.7+ and is built on top of the general-purpose simulation analysis package MDTraj^41^. SOURSOP uses MDTraj as a backend for parsing simulation trajectories and can accept trajectories in a wide variety of file formats. Although trajectory files are parsed into SOURSOP-specific objects, the underlying `mdtraj.topology` and `mdtraj.trajectory` objects remain user-facing and accessible. In this way, any analysis written to work with MDTraj is directly applicable to SOURSOP objects.

SOURSOP reads a simulation trajectory into a `SSTraj` object. The `SSTraj` object automatically extracts individual protein chains into their `SSProtein` objects. `SSProtein` objects are the base object upon which single-chain analysis routines are applied as object functions. In addition, peripheral modules that include `ssnmr` and `sspre`, provide modular, protein-independent analyses that work in conjunction with an `SSProtein` object. In this way, SOURSOP abides by the software principle of loose coupling, facilitating maintainability and future extension. The overall architecture of SOURSOP is shown in **Fig. 1A**.

**Figure 1:**
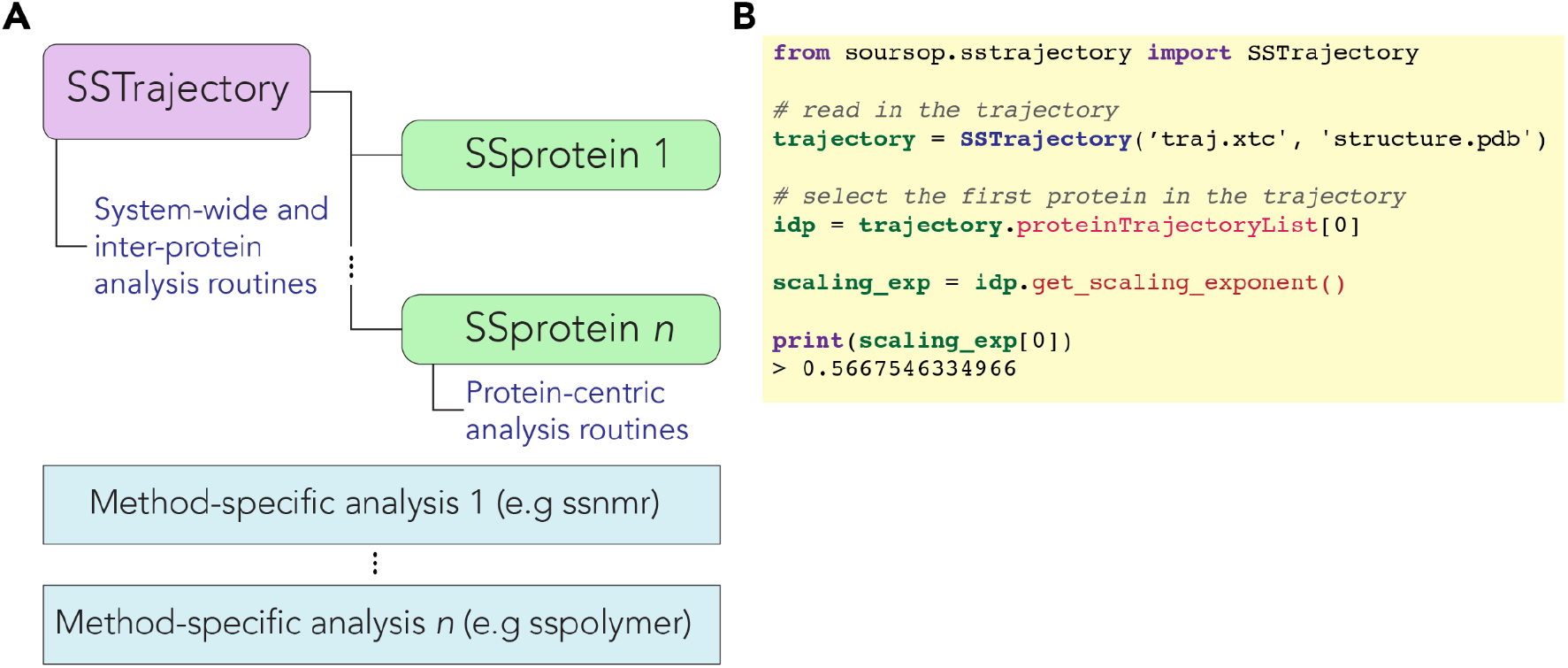
Architecture and example code for SOURSOP. **(A)** Trajectory files are read into an SSTrajectory object, which automatically parses each polypeptide chain into SSProtein objects. Each SSProtein object has a set of object-based analyses associated with them. Each trajectory must have between 1 and *n* protein chains in it. In addition, various stateless method-specific analysis modules exist for certain types of analysis. Additional stateless methods can be extended to allow new analysis routines to be incorporated in a way that does not alter the SSProtein or SSTrajectory code. **(B)** Example code illustrating how the apparent scaling exponent can be calculated from an ensemble.

Where possible and appropriate, SOURSOP engages in memoization, a dynamic programming approach where expensive calculations are saved after being executed once^52^. This offers a general strategy that avoids repeated recalculation of (for example) the same sets of distances. In addition to intramolecular analysis codified in the `SSProtein` object, intermolecular and multi-chain analysis routines are included in the `SSTraj` object. In this way, a simple and standardized interface for working with protein ensemble data is provided. Ensembles to be analyzed could be generated through standard all-atom simulations, but PDB ensembles from NMR or ensemble selection procedures are also directly analyzable.

A major goal in developing SOURSOP is to make simulation analysis easy and intuitive, both for the user and developers. As an example, **Fig. 1B** offers a simple example of computing the apparent scaling exponent (*ν*^app^) for a protein in a simulation trajectory. While a straightforward user experience is an obvious goal for any software package, providing a consistent, well-defined, and accessible software architecture is essential for long-term maintenance and extendibility. Well-structured software is also necessary to enable productive and sustainable open-source contributions.

The current working version can be found at https://github.com/holehouse-lab/soursop, with documentation at https://soursop.readthedocs.io/. SOURSOP uses PyTest (https://docs.pytest.org/en/stable/) for unit testing, Sphinx (https://www.sphinx-doc.org/en/master/), and readthedocs (https://readthedocs.org/) for documentation, and Git (https://git-scm.com/) and GitHub (https://github.com/) for version control. The original repository structure was generated using cookiecutter (https://github.com/cookiecutter/cookiecutter). Explicit dependencies include MDTraj^41^, SciPy^53^, NumPy^54^, Pandas^55^, and Cython^56^. In addition to the analyses shown here, we provide a collection of Jupyter notebooks along with the full trajectories (where possible) that offer examples of more general IDR-centric analysis that can be performed on the ensembles studied here (https://github.com/holehouse-lab/supportingdata/tree/master/2023/lalmansingh_2023). The SOURSOP code is consistent and heavily commented. The documentation also provides specific guidance for the development and integration of new analysis routines into SOURSOP.

## 3. RESULTS

To demonstrate the analyses available in SOURSOP, we have analyzed a collection of ensembles generated by various methods. The trajectories analyzed were generated using CAMPARI (an all-atom Monte Carlo simulation engine) or Desmond (an all-atom MD simulation engine)^26,57,58^. The analyses performed here are offered as convenient examples of the types of analyses and insight enabled by SOURSOP.

### 3.1 IDR global dimensions show extensive sequence-dependent conformational biases

A challenge in the study of IDRs is the absence of an obvious reference state. While folded proteins are typically associated with a native conformation which can serve as a reference point for further analysis, the structural heterogeneity of an IDR means that no single state serves this purpose. Conveniently, polymer physics offers analytical tools that can serve as reference states for disordered and unfolded protein ensembles ^30,48,59–63^. As a result, dimensionless polymeric parameters can be computed, which allows the conformational behavior of very different proteins to be quantitatively and directly compared. SOURSOP implements the calculation of many of these parameters, facilitating ensemble analysis.

We re-analyzed a series of conformational ensembles using two such dimensionless reference parameters. Specifically, we computed instantaneous asphericity (δ^*\**^), which measures the shape of a given conformation, and *t*, a dimensionless parameter that quantifies global dimensions effectively via a normalized radius of gyration as originally defined by Vitalis and Pappu as

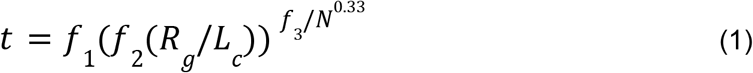

where *N* is the number of residues in the sequence, *L*_*c*_ is the contour length of the polypeptide in Angstroms (3.6×*N*), and *f*_*1*_, *f*_*2*_, and *f*_*3*_, are parameters used to ensure *t* remains in the interval of 0 to 1 and are defined as 2.5, 1.75 and 4.0. By generating 2D density plots that report on the simultaneous evaluation of *δ*^*\**^ and *t* for each conformation, a quantitative and length-normalized representation of IDR global conformational preferences can be easily visualized in a normalized manner. Both *t* and *δ*^*\**^ are transformations of the eigenvalues from the gyration tensor `T`. They represent global order parameters to describe the size and shape of a given conformation. An alternative normalization approach is using standard polymer models as reference states. To illustrate these two ideas, we use both approaches in this study.

We analyzed large conformational ensembles with over 3×10^4^ distinct conformers obtained from previously published simulations that have been directly benchmarked against experiments to compare how ensemble size and shape vary across different IDRs (**Fig. 2A, Table S1**) ^26,57,58,64–67^. This analysis revealed a wide array of global conformational behavior, with IDRs ranging from heterogeneous compact ensembles to highly expanded self-avoiding random chains commensurate with polypeptides under strongly denaturing conditions. To contextualize these global dimensions, we also calculated normalized radii of gyration using the dimensions of a sequence-matched chain under conditions in which chain-chain and chain-solvent interaction are counterbalanced, with similar results (**Fig. 2B**).

**Figure 2:**
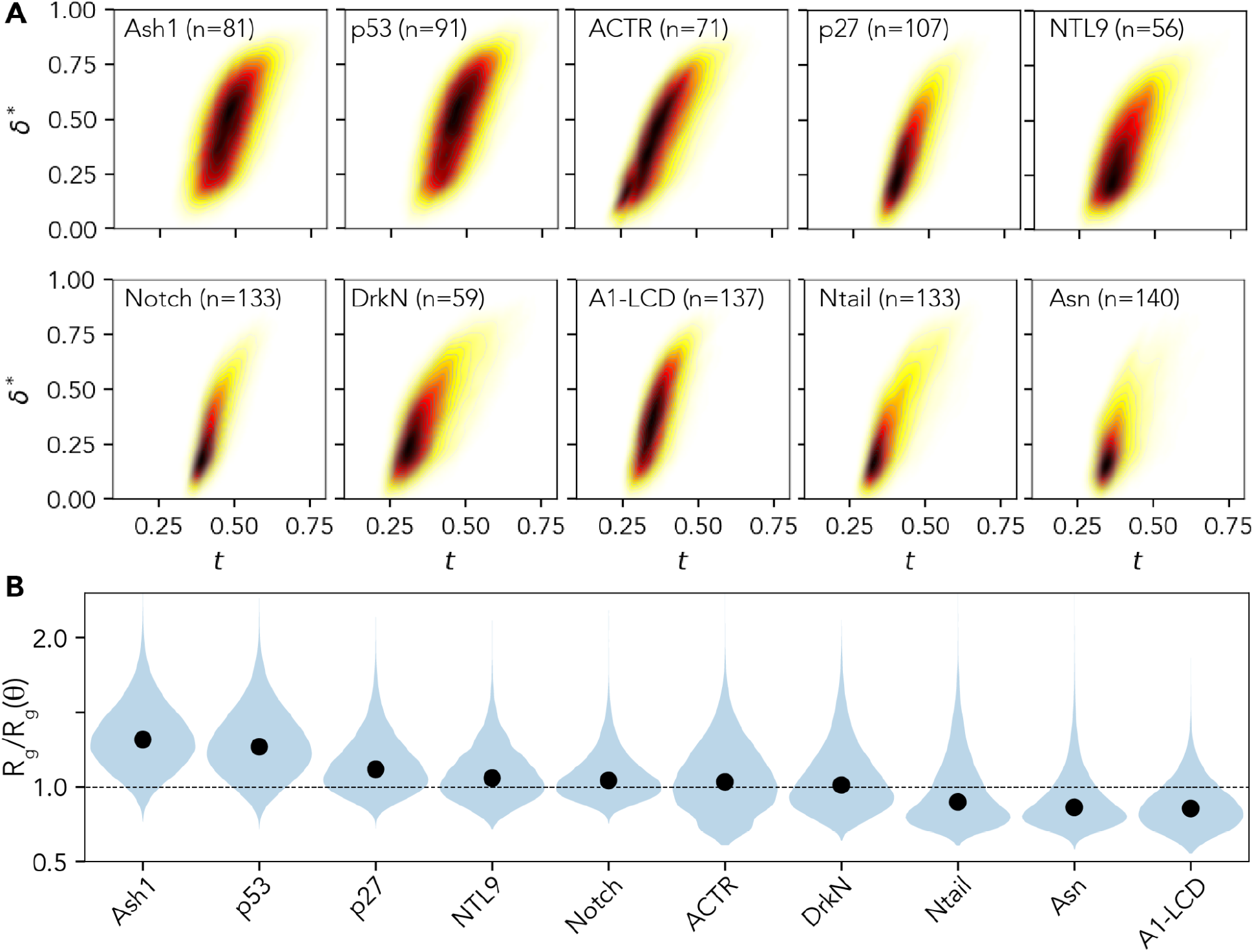
Global conformational analysis of 10 disordered protein ensembles analyzed with SOURSOP. **(A)** The two-dimensional density plots for instantaneous asphericity (δ*) and normalized dimensions (*t*) reveal a broad range of conformational landscapes. Ash1^58^, p53^68^, p27^67^, NTL9^66^, Notch^64^, and A1-LCD^57^ are ensembles generated by Monte Carlo ensembles with the ABSINTH implicit solvent model^60^. ACTR, drkN, NTail, and Asn are ensembles generated by molecular dynamics simulations with Amber99-disp forcefield^26^. Note that NTL9 is not an IDP, but the ensemble reported here represents an unfolded-state ensemble obtained under native conditions^66^. **(B)** Normalized chain dimensions were calculated by normalizing the instantaneous radius of gyration from ensembles by the expected radius of gyration from a sequence-matched chain in the theta state, whereby chain-chain and chain-solvent interactions are counterbalanced ^6,62,68^.

The diversity in global IDR properties (size and shape), as illustrated in **Fig. 2A**, is often masked by ensemble average properties. As a result, two IDRs may appear, on average, to be highly similar. The simulation analysis uncovers differences using the full distribution of conformations, which is evident even for relatively simple order parameters such as δ^*\**^ and *t*, in agreement with prior work that has shown ensemble-average properties can mask complexities in the underlying conformational ensemble ^69–71^.

### Aromatic and charged residues play an outsized role in dictating the conformational behavior of disordered proteins

Next, we applied SOURSOP to assess the sequence determinants of the attractive and repulsive intramolecular interactions that determine global and local conformational biases in IDR ensembles. To evaluate local chain interactions, we computed the radius of gyration over a sliding window of 14 residues to generate a linear profile of local density, normalizing for steric effects via an atomistic excluded volume (EV) model (**Fig. 3**, see **Supplemental Information**). To assess long-range interactions, we computed scaling maps (**Fig. 4**). Scaling maps are inter-residue distances normalized by the expected distances from some reference polymer model, in this case, the EV model. The use of scaling maps accounts for the intrinsic contribution that chain connectivity has to inter-residue distances. Across the set of ensembles examined, charged and aromatic residues emerge as key determinants of IDR local and global interactions irrespective of the forcefield or simulations approach being used (**Fig. 3, Fig. 4**).

**Figure 3:**
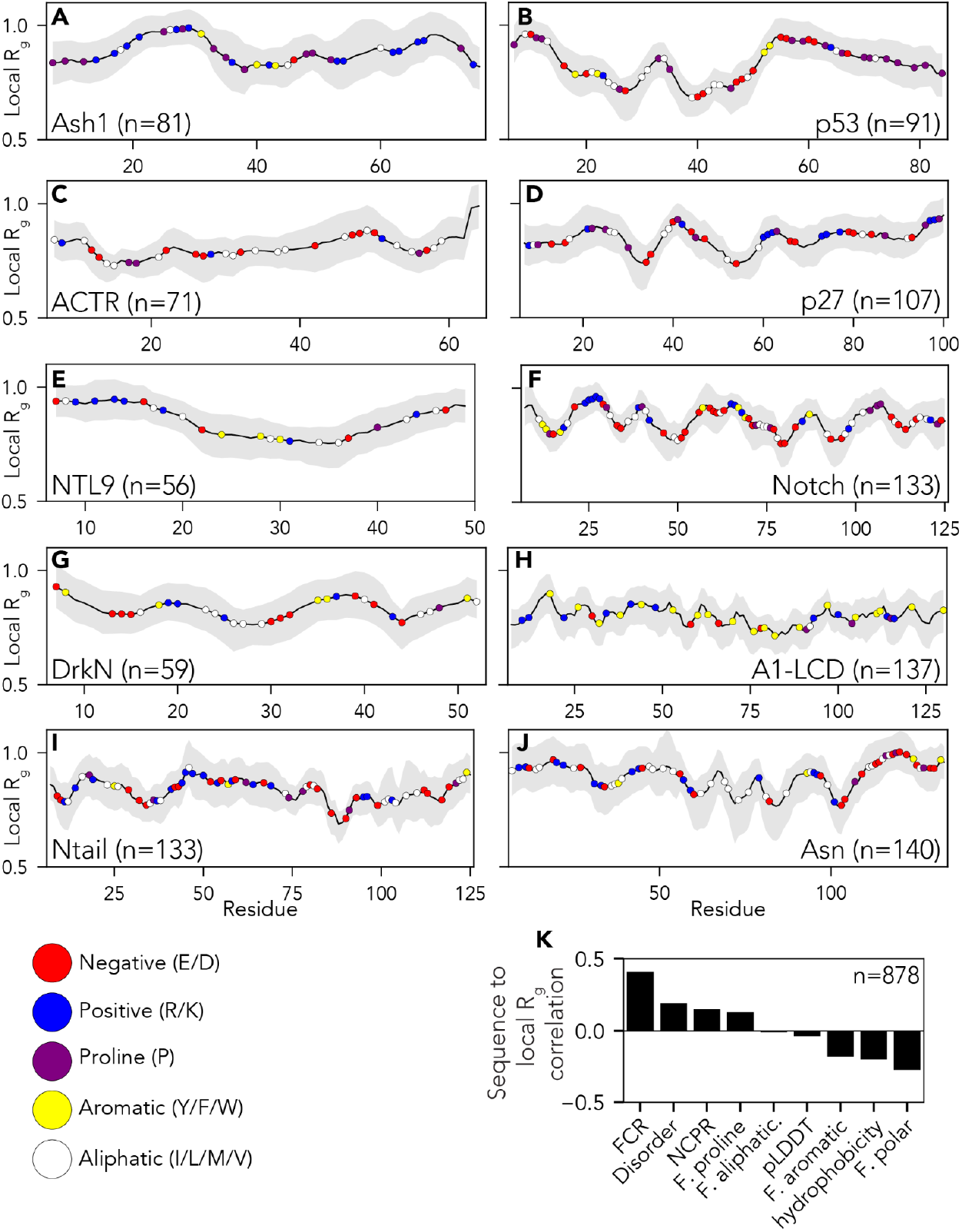
Local chain compaction with residue chemistry superimposed over the local radius of gyration (R_g_). (**A-J**) Individual plots showing analysis for each protein ensemble as introduced in Figure 2. Local R_g_ is calculated using a 14-residue sliding window. Colored circles on each plot represent different amino acid chemistry groups, highlighted in the legend below panel **I. (K)** Pearson’s correlation coefficient between local R_g_ and the amino acid chemistry within the window in question. The fraction of charged residues (FCR) is the strongest positive determinant of expansion and is stronger in these sequences than the net charge per residue (NCPR). While polar residues, in principle, correlate as negative determinants of expansion, the negative correlation is driven by subregions deficient in charged residues and enriched in only polar residues.

**Figure 4:**
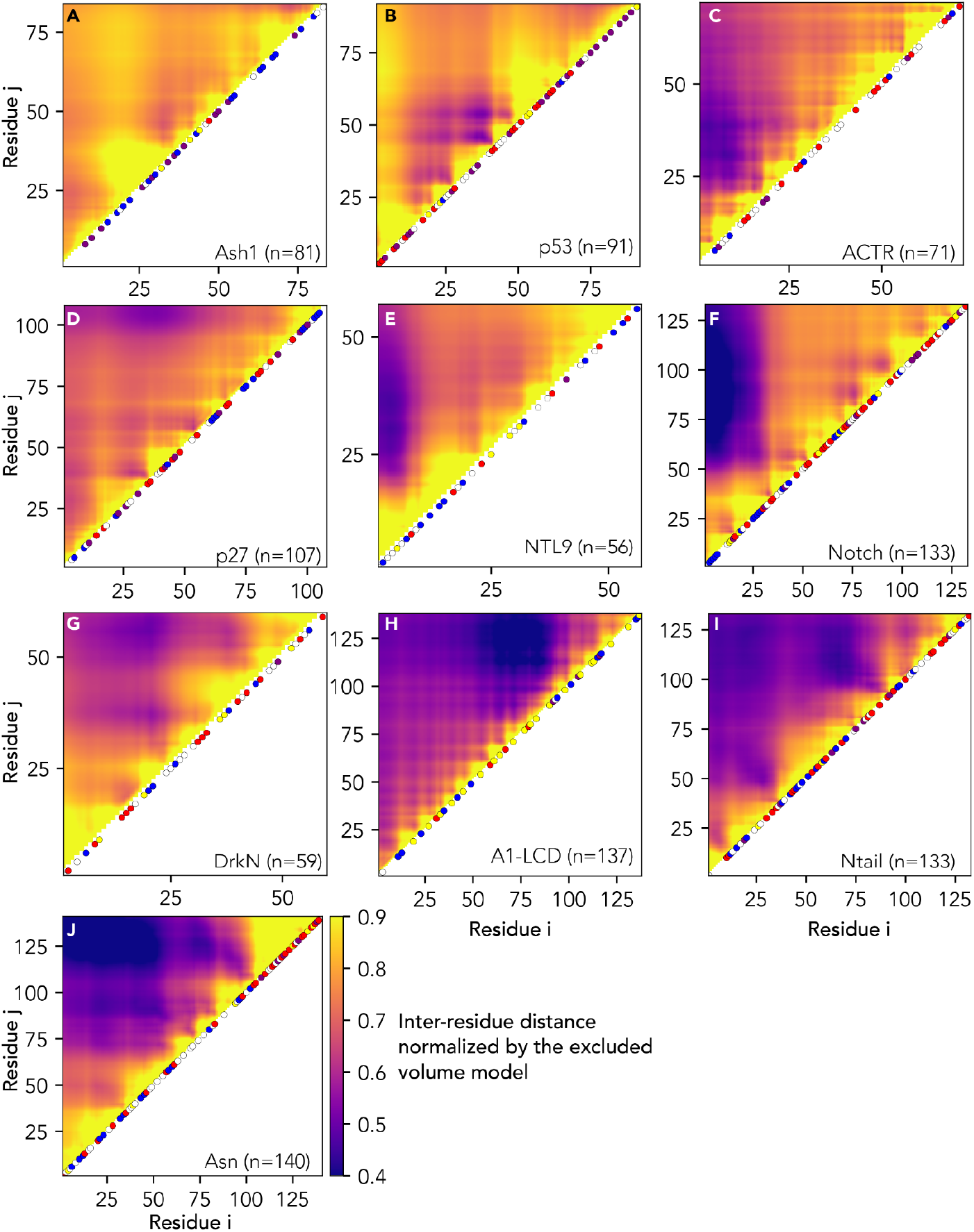
Preferential attraction and repulsion quantified via scaling maps that report the normalized distance between every pair of residues in the protein. (**A-J**) Individual plots of analysis for each protein ensemble as introduced in Figure 2. Normalized distances are calculated by dividing ensemble-average inter-residue distance by the distance obtained for the EV model. Attractive interactions emerge as darker colors, while repulsive interactions are lighter. Along the diagonal, subsets of residues are colored using the same color scheme used in Fig. 3.

While our analysis is necessarily retrospective and correlative, it is in line with prior experimental work^57,72–75^. To explore this observation further, we performed all-atom simulations using the ABSINTH implicit solvent model of the p53^1-91^ with three phosphomimetic mutations (S15E, T18E, S20E) and compared the result to previous simulations of the wildtype sequence (**Fig. 5A**)^65^. While glutamic acid is an imperfect analog for the larger phosphate group, the results revealed that relatively modest changes in linear charge density can cause local and long-range changes in the conformational ensemble. Despite substantial local conformational rearrangement, this leads only to a modest change of 0.5 Å in the mean radius of gyration (**Fig. 5B**). Charge effects leading to seemingly minor changes in global dimensions while altering local networks of intramolecular interactions mirrors prior work on the multi-phosphorylated proteins Ash1, Sic1, and a region of the RNA polymerase CTD ^58,76–78^. Taken together, these results suggest that while local changes in charge density can induce local conformational changes in ensemble behavior, compensatory changes in attractive (and repulsive) interactions that act on different or overlapping length scales can mask the effects of large-scale changes when global chain dimensions are examined.

**Figure 5:**
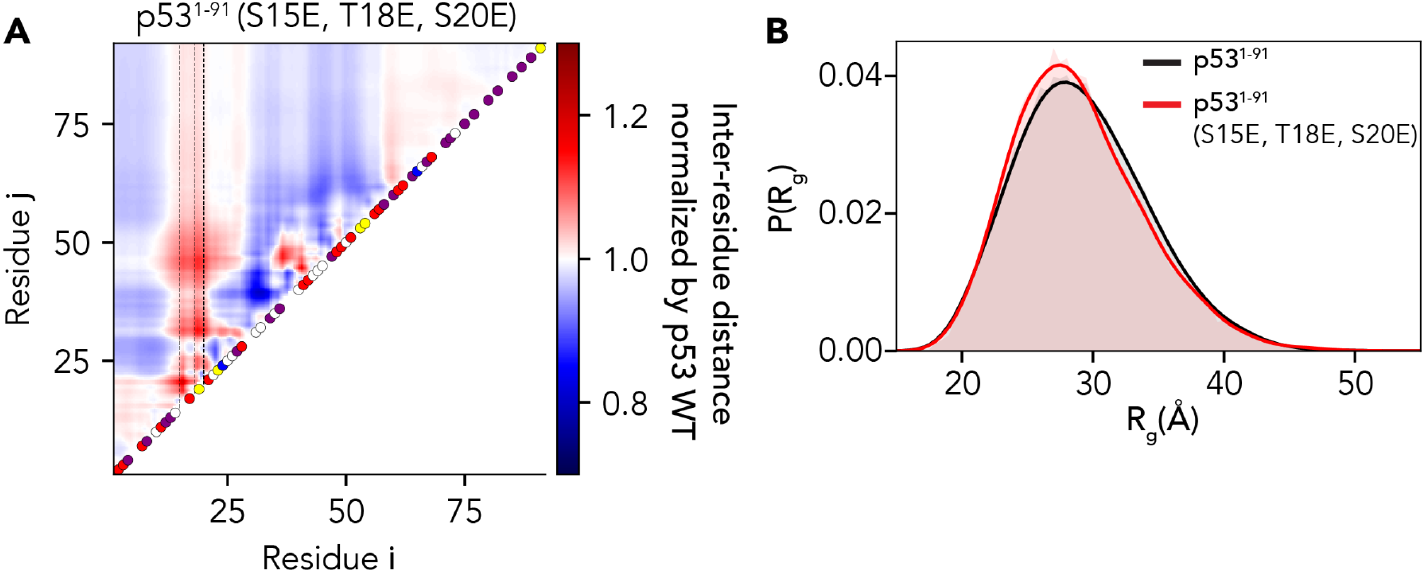
Comparison of changes in local and global dimensions for wildtype vs. phosphomimetic versions of p53. **(A)** Scaling maps where inter-residue distances for the phosphomimetic version of p53 N-terminal domain (p53^1-91^) are normalized by distances for the wild-type protein. Despite differing by only three residues in the N-terminal quarter of the protein, the phosphomimetic version of p53 shows substantial differences in long-range and local dimensions, as shown by the emergence of both attractive (blue) and repulsive (red) interactions. **(B)** Despite these rearrangements, a relatively small change in overall global dimensions is observed. While the wildtype ensemble-average R_g_ is 29.4 Å, the phosphomimetic variant is 29.1 Å, a difference below the statistical detection limits for most experimental techniques.

### Molecular accessibility is context dependent in IDRs

It is often convenient to imagine IDRs as uniformly accessible unfolded polypeptides. Under this model, each residue is equivalently solvent-accessible, and IDRs can be thought of as flexible scaffolds where the relative position along the chain has no real impact on molecular accessibility. While this is an appealingly simple model, given the complex conformational behavior observed in our analyses here and elsewhere, it may not be a given that every residue is equally accessible^58,70,79–82^. To examine this idea further, we computed local accessibility across eight-residue windows for each IDR using a 10 Å spherical probe (**Fig. 6**). This analysis allows us to assess how accessibility varies as a function of local sequence position.

**Figure 6:**
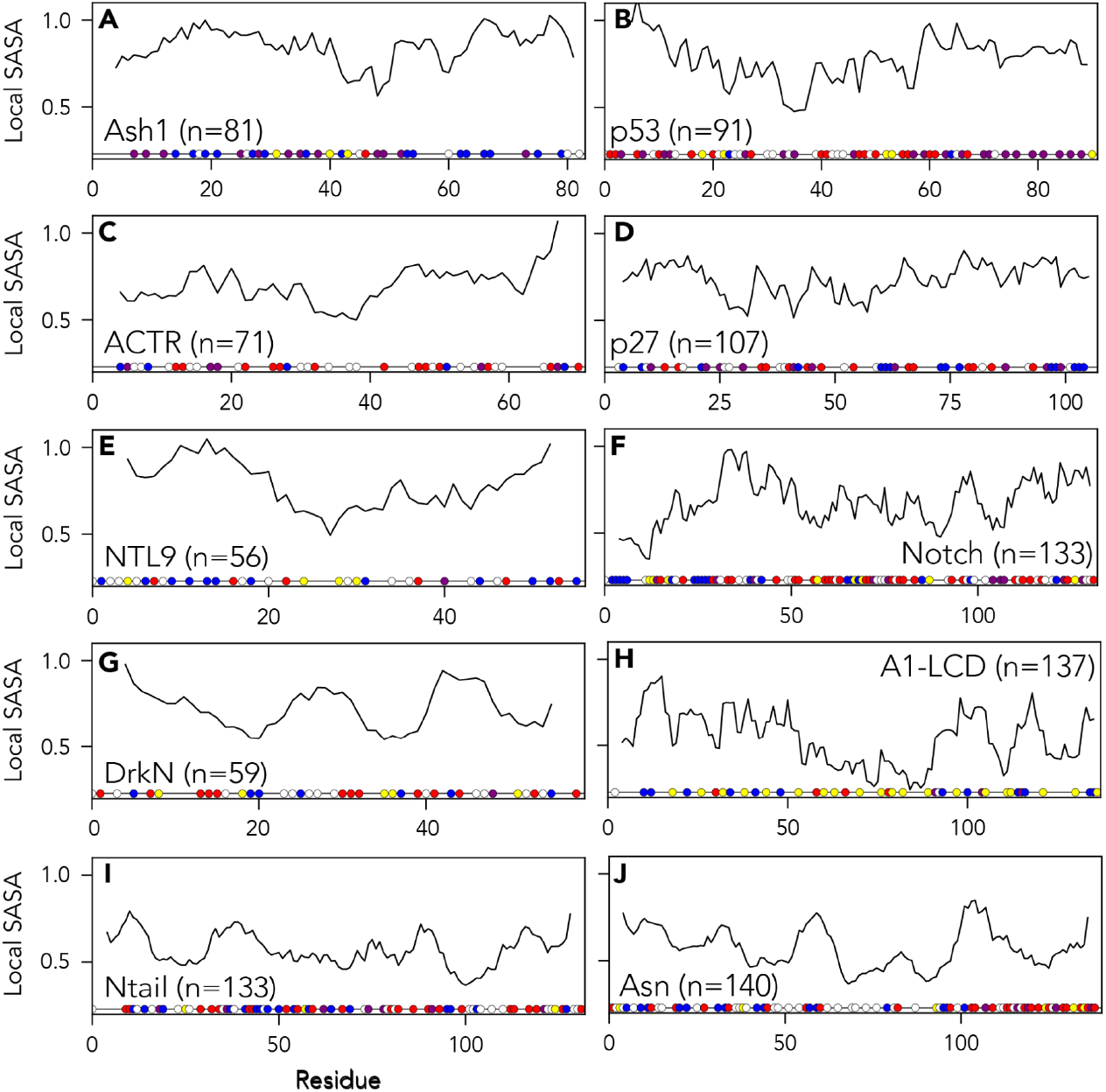
Normalized local solvent-accessible surface area (SASA) using an eight-residue sliding window and a 10 Å probe size. Normalization is done using excluded volume (EV) reference simulations to account for side-chain-dependent differences in solvent accessibility. Amino acid residues are colored as in Fig 3. Distinct patterns of accessibility are observed across different proteins, indicating long- and short-range intramolecular interactions can influence the accessibility of local binding sites.

Our analysis reveals substantial variation in molecular accessibility, suggesting that two residues of the same type may be differentially accessible depending on their broader sequence context (**Fig. 6**). Under this interpretation, the local sequence environment offers a putative mechanism to control the effective concentration of a local binding motif. The importance of local sequence context on molecular interactions can be further expanded if sequence-encoded chemistry provides partner-specific attractive and repulsive interactions. Taken together, despite the lack of a fixed 3D structure, it seems reasonable to speculate that the binding of motifs from IDRs should be considered both in terms of molecular sterics and shape complementarity (as is the conventional view for rigid-body molecular recognition) but also in terms of if and how the local chain context influences their accessibility and chemical context ^83,84^.

## 4. DISCUSSION

Here we introduce SOURSOP, an integrative Python-based software package for the analysis of all-atom ensembles extracted from simulations of intrinsically disordered proteins. SOURSOP is easy to install and use and is accompanied by extensive documentation and unit tests. Here we have shown how SOURSOP can be applied to analyze all-atom ensembles extracted from two types of simulations (Monte Carlo simulations and molecular dynamic simulations) of different IDRs. SOURSOP contains a range of additional routines not explored in this work but have been applied to various systems under a range of contexts, including local residual structure, intra-residue contacts, and the interaction between folded and disordered regions (**Fig. S1**)^57,58,85–87^.

### SOURSOP as a stand-alone package

SOURSOP was developed as a stand-alone analysis package built on the existing general-purpose simulation analysis package MDTraj^41^. The decision to develop SOURSOP as an independent package, as opposed to expanding the functionality of MDTraj, was motivated by several factors.

First, many of the analysis routines built into SOURSOP are of limited value for the analysis of well-folded proteins. At this juncture, MDTraj is a stable and mature software package that functions as the backend to a range of tools associated with molecular simulations ^44,88–92^. To add additional features into MDTraj would unavoidably lead to additional technical debt - more features to keep track of, manage, and test for. Technical debt adds viscosity, risks the introduction of new bugs, and can hamper future development if several coding styles are combined^93^. Accordingly, the drawbacks of integrating the analysis routines into MDTraj were judged to be substantially greater than the possible benefits.

Second, our goal is for SOURSOP to provide a general platform where novel analysis routines appropriate for disordered proteins can be implemented by the burgeoning community of labs performing simulations of disordered proteins. This requires our ability to maintain control over a consistent programmatic interface, which can be achieved via an interface layer between MDTraj and SOURSOP, but becomes challenging if analysis routines are implemented directly inside of MDTraj. For this reason, providing SOURSOP as a loosely-coupled software component that works with MDTraj, as opposed to within MDTraj, enables the best of both worlds.

Finally, applying principles from polymer physics to analyze disordered proteins is not new. Several of the analysis routines provided by SOURSOP are also available in extant software, notably in the simulation engine CAMPARI (http://campari.sourceforge.net/) ^6,30,60,61,94^. SOURSOP provides a lightweight toolkit that is simple to install, simple to use, and interoperable with MDTraj and the collection of existing analysis tools therein. Therefore, while some overlap exists, we do not see SOURSOP as replacing the analysis routines in MDTraj or CAMPARI. Instead, SOURSOP is a complement to extant routines and packages. Furthermore, it makes it relatively straightforward for groups to publish scripts or Jupyter notebooks that enable full reproduction of their analysis workflow.

### SOURSOP in the broader ecosystem of simulation software

Over the last three decades, considerable effort has been placed into the development of Monte Carlo (MC) and Molecular Dynamics (MD) methods for the accurate modeling of proteins and related systems *in silico*^*26–33*^. As advancements in computational resources emerged, they enabled the increasingly detailed estimations and characterizations of biophysical properties that would ordinarily be inaccessible through traditional experimental methods via the development of new modeling techniques and simulation paradigms.

As the simulation tools matured, new representational paradigms that enabled the simulation of systems at longer timescales also emerged beyond all-atom methods. Of note are two methods that provide fast results with reasonable accuracy: united-atom modeling and coarse-graining. United-atom modeling represents groups of molecular atoms as a particle. Coarse-grained representations allow groups of atoms, individual residues, or collections of residues to be represented as a particle. Similar to representational encoding, specific toolkits were developed for modeling specific system types and sizes, including lipids, membrane proteins, *etc*. Consequently, molecular modeling has become an important approach for understanding protein-protein interactions, protein folding, drug design, ligand-binding affinities, and much more.

However, as the timescale of such simulations increases with improvements in hardware, elucidating biological insights from the resultant trajectories becomes more difficult due to the requirement of more computational resources and time. These limiting factors resulted in the emergence of numerous analysis tools and frameworks to efficiently extract various attributes of interest based on the system type and size. Properties include the radius of gyration, Principal Component Analysis (PCA) of structural properties, cluster analysis, Root Mean Square Distance (RMSD) calculations, Root Mean Square Fluctuations (RMSF), Radial Distribution Function calculations, and many more. An extended discussion of the current ecosystem of simulation analysis packages is provided in the **Supplemental Information**.

### SOURSOP is an extendable platform for novel analysis routines

Analyzing IDR ensembles to reveal clear and interpretable conclusions remains challenging. Absent a native reference state, it can be difficult to generate informative and visually coherent representations that fully capture the inherent high dimensionality of an IDR ensemble. While various ‘standard’ analyses have emerged for folded proteins (*e.g*., contact maps, per-residue RMSF, the fraction of native contacts), there is less consensus on what the standard analyses should be when assessing IDR ensembles.

Rather than a problem, this raises an opportunity for innovation, whereby novel analysis and visualization approaches are needed. With this in mind, we hope new analysis routines can be integrated into SOURSOP, facilitating distribution and packaging. Considering this objective, SOURSOP includes a well-defined style guide for new analysis routines and a collection of utility functions that provide automatic sanity checking and defensive programming for input data. We also provide documentation on how best to introduce a new routine and how to integrate it into the main codebase. These features, combined with the broad reach of the Python programming language, will lower the barrier to open-source and community-driven scientific development.

## 5. CONCLUSION

SOURSOP is an open-source Python toolkit for the general analysis of ensembles of disordered proteins. In addition to analyzing disordered protein ensembles, SOURSOP can also be used to analyze folded protein trajectories or individual PDB files. As such, SOURSOP offers a general interface for calculating molecular properties, polymeric parameters, and the development of new IDR-centric analysis routines.

## ACKNOWLEDGMENTS

We thank all members of the Pappu and Holehouse labs for extensive testing, error reporting, and feature requests in SOURSOP. The development of SOURSOP was financially and intellectually supported by the Molecular Sciences Software Institute (MOLSSI), and A.S.H. received an NSF-sponsored MOLSSI postdoctoral fellowship (NSF-1547580 (subaward 479590)). This work was also supported by NSF grant no. 2128068 to ASH, and we thank members of the Water and Life Interface Institute (WALII), supported by NSF DBI grant #2213983, for helpful discussions. J.M.L. is a data scientist and scientific programmer in the Center for Biomolecular Condensates at Washington University in St. Louis. Work by members of the Pappu lab was supported by a grant from the Air Force Office of Scientific Research (FA9550-20-1-0241) and the US National Institutes of Health (R01NS121114).

SOURSOP was previously called CTraj and CAMPARITraj. We renamed the package SOURSOP to avoid confusion and the incorrect implication that this is a CAMPARI-specific analysis package.

## Notes

https://github.com/holehouse-lab/soursop

https://soursop.readthedocs.io/

### Competing Interest Statement

The authors have declared no competing interest.

## REFERENCES

(1) van der Lee, R.; Buljan, M.; Lang, B.; Weatheritt, R. J.; Daughdrill, G. W.; Dunker, A. K.; Fuxreiter, M.; Gough, J.; Gsponer, J.; Jones, D. T.; Kim, P. M.; Kriwacki, R. W.; Oldfield, C. J.; Pappu, R. V.; Tompa, P.; Uversky, V. N.; Wright, P. E.; Babu, M. M. Classification of Intrinsically Disordered Regions and Proteins. Chem. Rev. 2014, 114 (13), 6589–6631.

(2) Sigler, P. B. Acid Blobs & Negative Noodles. Nature 1988, 333, 210–212.

(3) Ptitsyn, O. B.; Uversky, V. N. The Molten Globule Is a Third Thermodynamical State of Protein Molecules. FEBS Lett. 1994, 341 (1), 15–18.

(4) Wright, P. E.; Dyson, H. J. Intrinsically Unstructured Proteins: Re-Assessing the Protein Structure-Function Paradigm. J. Mol. Biol. 1999, 293 (2), 321–331.

(5) Babu, M. M.; Kriwacki, R. W.; Pappu, R. V. Structural Biology. Versatility from Protein Disorder. Science 2012, 337 (6101), 1460–1461.

(6) Mao, A. H.; Lyle, N.; Pappu, R. V. Describing Sequence-Ensemble Relationships for Intrinsically Disordered Proteins. Biochem. J 2013, 449 (2), 307–318.

(7) Dyson, H. J.; Wright, P. E. Unfolded Proteins and Protein Folding Studied by NMR. Chem. Rev. 2004, 104 (8), 3607–3622.

(8) Mittag, T.; Forman-Kay, J. D. Atomic-Level Characterization of Disordered Protein Ensembles. Curr. Opin. Struct. Biol. 2007, 17 (1), 3–14.

(9) Das, R. K.; Ruff, K. M.; Pappu, R. V. Relating Sequence Encoded Information to Form and Function of Intrinsically Disordered Proteins. Curr. Opin. Struct. Biol. 2015, 32 (0), 102–112.

(10) Martin, E. W.; Holehouse, A. S. Intrinsically Disordered Protein Regions and Phase Separation: Sequence Determinants of Assembly or Lack Thereof. Emerg Top Life Sci 2020, 4 (3), 307–329.

(11) Schuler, B.; Soranno, A.; Hofmann, H.; Nettels, D. Single-Molecule FRET Spectroscopy and the Polymer Physics of Unfolded and Intrinsically Disordered Proteins. Annu. Rev. Biophys. 2016, 45, 207–231.

(12) Kikhney, A. G.; Svergun, D. I. A Practical Guide to Small Angle X-ray Scattering (SAXS) of Flexible and Intrinsically Disordered Proteins. FEBS Lett. 2015.

(13) Chemes, L. B.; Alonso, L. G.; Noval, M. G.; de Prat-Gay, G. Circular Dichroism Techniques for the Analysis of Intrinsically Disordered Proteins and Domains. Methods Mol. Biol. 2012, 895, 387–404.

(14) Gast, K.; Fiedler, C. Dynamic and Static Light Scattering of Intrinsically Disordered Proteins. Methods Mol. Biol. 2012, 896, 137–161.

(15) Jensen, M. R.; Zweckstetter, M.; Huang, J.-R.; Blackledge, M. Exploring Free-Energy Landscapes of Intrinsically Disordered Proteins at Atomic Resolution Using NMR Spectroscopy. Chem. Rev. 2014, 114 (13), 6632–6660.

(16) Gibbs, E. B.; Cook, E. C.; Showalter, S. A. Application of NMR to Studies of Intrinsically Disordered Proteins. Arch. Biochem. Biophys. 2017, 628, 57–70.

(17) Best, R. B. Computational and Theoretical Advances in Studies of Intrinsically Disordered Proteins. Curr. Opin. Struct. Biol. 2017, 42, 147–154.

(18) Wang, X.; Vitalis, A.; Wyczalkowski, M. A.; Pappu, R. V. Characterizing the Conformational Ensemble of Monomeric Polyglutamine. Proteins 2006, 63 (2), 297–311.

(19) Shea, J.-E.; Best, R. B.; Mittal, J. Physics-Based Computational and Theoretical Approaches to Intrinsically Disordered Proteins. Curr. Opin. Struct. Biol. 2021, 67, 219–225.

(20) Alston, J. J.; Soranno, A.; Holehouse, A. S. Integrating Single-Molecule Spectroscopy and Simulations for the Study of Intrinsically Disordered Proteins. Methods 2021, 193, 116–135.

(21) Palazzesi, F.; Prakash, M. K.; Bonomi, M.; Barducci, A. Accuracy of Current All-Atom Force-Fields in Modeling Protein Disordered States. J. Chem. Theory Comput. 2015, 11 (1), 2–7.

(22) Zerze, G. H.; Zheng, W.; Best, R. B.; Mittal, J. Evolution of All-Atom Protein Force Fields to Improve Local and Global Properties. J. Phys. Chem. Lett. 2019, 10 (9), 2227–2234.

(23) Ruff, K. M.; Pappu, R. V.; Holehouse, A. S. Conformational Preferences and Phase Behavior of Intrinsically Disordered Low Complexity Sequences: Insights from Multiscale Simulations. Curr. Opin. Struct. Biol. 2019, 56, 1–10.

(24) Karplus, M.; Petsko, G. A. Molecular Dynamics Simulations in Biology. Nature 1990, 347 (6294), 631–639.

(25) Bottaro, S.; Lindorff-Larsen, K. Biophysical Experiments and Biomolecular Simulations: A Perfect Match? Science 2018, 361 (6400), 355–360.

(26) Robustelli, P.; Piana, S.; Shaw, D. E. Developing a Molecular Dynamics Force Field for Both Folded and Disordered Protein States. Proc. Natl. Acad. Sci. U. S. A. 2018, 115 (21), E4758–E4766.

(27) Piana, S.; Robustelli, P.; Tan, D.; Chen, S.; Shaw, D. E. Development of a Force Field for the Simulation of Single-Chain Proteins and Protein-Protein Complexes. J. Chem. Theory Comput. 2020, 16 (4), 2494–2507.

(28) Best, R. B.; Mittal, J. Protein Simulations with an Optimized Water Model: Cooperative Helix Formation and Temperature-Induced Unfolded State Collapse. J. Phys. Chem. B 2010, 114 (46), 14916–14923.

(29) Best, R. B.; Zheng, W.; Mittal, J. Balanced Protein-Water Interactions Improve Properties of Disordered Proteins and Non-Specific Protein Association. J. Chem. Theory Comput. 2014, 10 (11), 5113–5124.

(30) Vitalis, A.; Pappu, R. V. ABSINTH: A New Continuum Solvation Model for Simulations of Polypeptides in Aqueous Solutions. J. Comput. Chem. 2009, 30 (5), 673–699.

(31) Huang, J.; Rauscher, S.; Nawrocki, G.; Ran, T.; Feig, M.; de Groot, B. L.; Grubmüller, H.; MacKerell, A. D., Jr. CHARMM36m: An Improved Force Field for Folded and Intrinsically Disordered Proteins. Nat. Methods 2017, 14 (1), 71–73.

(32) Piana, S.; Donchev, A. G.; Robustelli, P.; Shaw, D. E. Water Dispersion Interactions Strongly Influence Simulated Structural Properties of Disordered Protein States. J. Phys. Chem. B 2015, 119 (16), 5113–5123.

(33) Appadurai, R.; Nagesh, J.; Srivastava, A. High Resolution Ensemble Description of Metamorphic and Intrinsically Disordered Proteins Using an Efficient Hybrid Parallel Tempering Scheme. Nat. Commun. 2021, 12 (1), 958.

(34) Salomon-Ferrer, R.; Case, D. A.; Walker, R. C. An Overview of the Amber Biomolecular Simulation Package. Wiley Interdiscip. Rev. Comput. Mol. Sci. 2013, 3 (2), 198–210.

(35) Brooks, B. R.; Brooks, C. L., 3rd; Mackerell, A. D., Jr; Nilsson, L.; Petrella, R. J.; Roux, B.; Won, Y.; Archontis, G.; Bartels, C.; Boresch, S.; Caflisch, A.; Caves, L.; Cui, Q.; Dinner, A. R.; Feig, M.; Fischer, S.; Gao, J.; Hodoscek, M.; Im, W.; Kuczera, K.; Lazaridis, T.; Ma, J.; Ovchinnikov, V.; Paci, E.; Pastor, R. W.; Post, C. B.; Pu, J. Z.; Schaefer, M.; Tidor, B.; Venable, R. M.; Woodcock, H. L.; Wu, X.; Yang, W.; York, D. M.; Karplus, M. CHARMM: The Biomolecular Simulation Program. J. Comput. Chem. 2009, 30 (10), 1545–1614.

(36) Abraham, M. J.; Murtola, T.; Schulz, R.; Páll, S.; Smith, J. C.; Hess, B.; Lindahl, E. GROMACS: High Performance Molecular Simulations through Multi-Level Parallelism from Laptops to Supercomputers. SoftwareX 2015/9, 1–2, 19–25.

(37) Eastman, P.; Swails, J.; Chodera, J. D.; McGibbon, R. T.; Zhao, Y.; Beauchamp, K. A.; Wang, L.-P.; Simmonett, A. C.; Harrigan, M. P.; Stern, C. D.; Wiewiora, R. P.; Brooks, B. R.; Pande, V. S. OpenMM 7: Rapid Development of High Performance Algorithms for Molecular Dynamics. PLoS Comput. Biol. 2017, 13 (7), e1005659.

(38) Thompson, A. P.; Aktulga, H. M.; Berger, R.; Bolintineanu, D. S.; Brown, W. M.; Crozier, P. S.; in ‘t Veld, P. J.; Kohlmeyer, A.; Moore, S. G.; Nguyen, T. D.; Shan, R.; Stevens, M. J.; Tranchida, J.; Trott, C.; Plimpton, S. J. LAMMPS - a Flexible Simulation Tool for Particle-Based Materials Modeling at the Atomic, Meso, and Continuum Scales. Comput. Phys. Commun. 2022, 271, 108171.

(39) Bowers, K. J.; Chow, E.; Xu, H.; Dror, R. O.; Eastwood, M. P.; Gregersen, B. A.; Klepeis, J. L.; Kolossvary, I.; Moraes, M. A.; Sacerdoti, F. D.; Salmon, J. K.; Shan, Y.; Shaw, D. E. Scalable Algorithms for Molecular Dynamics Simulations on Commodity Clusters. In Proceedings of the 2006 ACM/IEEE conference on Supercomputing; SC ‘06; Association for Computing Machinery: New York, NY, USA, 2006; p 84–es.

(40) Phillips, J. C.; Hardy, D. J.; Maia, J. D. C.; Stone, J. E.; Ribeiro, J. V.; Bernardi, R. C.; Buch, R.; Fiorin, G.; Hénin, J.; Jiang, W.; McGreevy, R.; Melo, M. C. R.; Radak, B. K.; Skeel, R. D.; Singharoy, A.; Wang, Y.; Roux, B.; Aksimentiev, A.; Luthey-Schulten, Z.; Kalé, L. V.; Schulten, K.; Chipot, C.; Tajkhorshid, E. Scalable Molecular Dynamics on CPU and GPU Architectures with NAMD. J. Chem. Phys. 2020, 153 (4), 044130.

(41) McGibbon, R. T.; Beauchamp, K. A.; Harrigan, M. P.; Klein, C.; Swails, J. M.; Hernández, C. X.; Schwantes, C. R.; Wang, L.-P.; Lane, T. J.; Pande, V. S. MDTraj: A Modern, Open Library for the Analysis of Molecular Dynamics Trajectories. Biophys. J. 2015, 109 (8), 1528–1532.

(42) Michaud-Agrawal, N.; Denning, E. J.; Woolf, T. B.; Beckstein, O. MDAnalysis: A Toolkit for the Analysis of Molecular Dynamics Simulations. J. Comput. Chem. 2011, 32 (10), 2319–2327.

(43) Romo, T. D.; Leioatts, N.; Grossfield, A. Lightweight Object Oriented Structure Analysis: Tools for Building Tools to Analyze Molecular Dynamics Simulations. J. Comput. Chem. 2014, 35 (32), 2305–2318.

(44) Porter, J. R.; Zimmerman, M. I.; Bowman, G. R. Enspara: Modeling Molecular Ensembles with Scalable Data Structures and Parallel Computing. J. Chem. Phys. 2019, 150 (4), 044108.

(45) Scherer, M. K.; Trendelkamp-Schroer, B.; Paul, F.; Pérez-Hernández, G.; Hoffmann, M.; Plattner, N.; Wehmeyer, C.; Prinz, J.-H.; Noé, F. PyEMMA 2: A Software Package for Estimation, Validation, and Analysis of Markov Models. J. Chem. Theory Comput. 2015, 11 (11), 5525–5542.

(46) Beauchamp, K. A.; Bowman, G. R.; Lane, T. J.; Maibaum, L.; Haque, I. S.; Pande, V. S. MSMBuilder2: Modeling Conformational Dynamics at the Picosecond to Millisecond Scale. J. Chem. Theory Comput. 2011, 7 (10), 3412–3419.

(47) Levine, Z. A.; Shea, J.-E. Simulations of Disordered Proteins and Systems with Conformational Heterogeneity. Curr. Opin. Struct. Biol. 2017, 43, 95–103.

(48) Pappu, R. V.; Wang, X.; Vitalis, A.; Crick, S. L. A Polymer Physics Perspective on Driving Forces and Mechanisms for Protein Aggregation - Highlight Issue: Protein Folding. Arch. Biochem. Biophys. 2008, 469 (1), 132–141.

(49) Hofmann, H.; Soranno, A.; Borgia, A.; Gast, K.; Nettels, D.; Schuler, B. Polymer Scaling Laws of Unfolded and Intrinsically Disordered Proteins Quantified with Single-Molecule Spectroscopy. Proc. Natl. Acad. Sci. U. S. A. 2012, 109 (40), 16155–16160.

(50) Cubuk, J.; Soranno, A. Macromolecular Crowding and Intrinsically Disordered Proteins: A Polymer Physics Perspective. ChemSystemsChem 2022. https://doi.org/10.1002/syst.202100051.

(51) Sørensen, C. S.; Kjaergaard, M. Effective Concentrations Enforced by Intrinsically Disordered Linkers Are Governed by Polymer Physics. Proc. Natl. Acad. Sci. U. S. A. 2019, 116 (46), 23124–23131.

(52) Cormen, T. H.; Leiserson, C. E.; Rivest, R. L.; Stein, C. Introduction to Algorithms, Fourth Edition; MIT Press, 2022.

(53) Virtanen, P.; Gommers, R.; Oliphant, T. E.; Haberland, M.; Reddy, T.; Cournapeau, D.; Burovski, E.; Peterson, P.; Weckesser, W.; Bright, J.; van der Walt, S. J.; Brett, M.; Wilson, J.; Millman, K. J.; Mayorov, N.; Nelson, A. R. J.; Jones, E.; Kern, R.; Larson, E.; Carey, C. J.; Polat, İ.; Feng, Y.; Moore, E. W.; VanderPlas, J.; Laxalde, D.; Perktold, J.; Cimrman, R.; Henriksen, I.; Quintero, E. A.; Harris, C. R.; Archibald, A. M.; Ribeiro, A. H.; Pedregosa, F.; van Mulbregt, P.; SciPy 1.0 Contributors. SciPy 1.0: Fundamental Algorithms for Scientific Computing in Python. Nat. Methods 2020, 17 (3), 261–272.

(54) van der Walt, S.; Colbert, S. C.; Varoquaux, G. The NumPy Array: A Structure for Efficient Numerical Computation. Computing in Science Engineering 2011, 13 (2), 22–30.

(55) McKinney, W. DAta STructures for STatistical COmputing in PYthon. In Proceedings of the 9th Python in Science Conference; SciPy, 2010. https://doi.org/10.25080/majora-92bf1922-00a.

(56) Behnel, S.; Bradshaw, R.; Citro, C.; Dalcin, L.; Seljebotn, D. S.; Smith, K. Cython: The Best of Both Worlds. Computing in Science Engineering 2011, 13 (2), 31–39.

(57) Martin, E. W.; Holehouse, A. S.; Peran, I.; Farag, M.; Incicco, J. J.; Bremer, A.; Grace, C. R.; Soranno, A.; Pappu, R. V.; Mittag, T. Valence and Patterning of Aromatic Residues Determine the Phase Behavior of Prion-like Domains. Science 2020, 367 (6478), 694–699.

(58) Martin, E. W.; Holehouse, A. S.; Grace, C. R.; Hughes, A.; Pappu, R. V.; Mittag, T. Sequence Determinants of the Conformational Properties of an Intrinsically Disordered Protein Prior to and upon Multisite Phosphorylation. J. Am. Chem. Soc. 2016, 138 (47), 15323–15335.

(59) Crick, S. L.; Pappu, R. V. Thermodynamic and Kinetic Models for Aggregation of Intrinsically Disordered Proteins. Protein and Peptide Folding, Misfolding, and Non-Folding 2012, 413–440.

(60) Vitalis, A.; Pappu, R. V. Chapter 3 Methods for Monte Carlo Simulations of Biomacromolecules. In Annual Reports in Computational Chemistry; Wheeler, R. A., Ed.; Elsevier, 2009; Vol. 5, pp 49–76.

(61) Vitalis, A.; Wang, X.; Pappu, R. V. Quantitative Characterization of Intrinsic Disorder in Polyglutamine: Insights from Analysis Based on Polymer Theories. Biophys. J. 2007, 93 (6), 1923–1937.

(62) Holehouse, A. S.; Garai, K.; Lyle, N.; Vitalis, A.; Pappu, R. V. Quantitative Assessments of the Distinct Contributions of Polypeptide Backbone Amides versus Side Chain Groups to Chain Expansion via Chemical Denaturation. J. Am. Chem. Soc. 2015, 137 (8), 2984–2995.

(63) Zerze, G. H.; Best, R. B.; Mittal, J. Sequence- and Temperature-Dependent Properties of Unfolded and Disordered Proteins from Atomistic Simulations. J. Phys. Chem. B 2015, 119 (46), 14622–14630.

(64) Sherry, K. P.; Das, R. K.; Pappu, R. V.; Barrick, D. Control of Transcriptional Activity by Design of Charge Patterning in the Intrinsically Disordered RAM Region of the Notch Receptor. Proc. Natl. Acad. Sci. U. S. A. 2017, 114 (44), E9243–E9252.

(65) Holehouse, A. S.; Sukenik, S. Controlling Structural Bias in Intrinsically Disordered Proteins Using Solution Space Scanning. J. Chem. Theory Comput. 2020, 16 (3), 1794–1805.

(66) Peran, I.; Holehouse, A. S.; Carrico, I. S.; Pappu, R. V.; Bilsel, O.; Raleigh, D. P. Unfolded States under Folding Conditions Accommodate Sequence-Specific Conformational Preferences with Random Coil-like Dimensions. Proc. Natl. Acad. Sci. U. S. A. 2019, 116 (25), 12301–12310.

(67) Das, R. K.; Huang, Y.; Phillips, A. H.; Kriwacki, R. W.; Pappu, R. V. Cryptic Sequence Features within the Disordered Protein p27Kip1 Regulate Cell Cycle Signaling. Proc. Natl. Acad. Sci. U. S. A. 2016, 113 (20), 5616–5621.

(68) Moses, D.; Yu, F.; Ginell, G. M.; Shamoon, N. M.; Koenig, P. S.; Holehouse, A. S.; Sukenik, S. Revealing the Hidden Sensitivity of Intrinsically Disordered Proteins to Their Chemical Environment. J. Phys. Chem. Lett. 2020, 11 (23), 10131–10136.

(69) Song, J.; Gomes, G.-N.; Shi, T.; Gradinaru, C. C.; Chan, H. S. Conformational Heterogeneity and FRET Data Interpretation for Dimensions of Unfolded Proteins. Biophys. J. 2017, 113 (5), 1012–1024.

(70) Fuertes, G.; Banterle, N.; Ruff, K. M.; Chowdhury, A.; Mercadante, D.; Koehler, C.; Kachala, M.; Estrada Girona, G.; Milles, S.; Mishra, A.; Onck, P. R.; Gräter, F.; Esteban-Martín, S.; Pappu, R. V.; Svergun, D. I.; Lemke, E. A. Decoupling of Size and Shape Fluctuations in Heteropolymeric Sequences Reconciles Discrepancies in SAXS vs. FRET Measurements. Proc. Natl. Acad. Sci. U. S. A. 2017, 114 (31), E6342–E6351.

(71) Ruff, K. M.; Holehouse, A. S. SAXS versus FRET: A Matter of Heterogeneity? Biophys. J. 2017, 113 (5), 971–973.

(72) Marsh, J. A.; Forman-Kay, J. D. Sequence Determinants of Compaction in Intrinsically Disordered Proteins. Biophys. J. 2010, 98 (10), 2383–2390.

(73) Mao, A. H.; Crick, S. L.; Vitalis, A.; Chicoine, C. L.; Pappu, R. V. Net Charge per Residue Modulates Conformational Ensembles of Intrinsically Disordered Proteins. Proc. Natl. Acad. Sci. U. S. A. 2010, 107 (18), 8183–8188.

(74) Müller-Späth, S.; Soranno, A.; Hirschfeld, V.; Hofmann, H.; Rüegger, S.; Reymond, L.; Nettels, D.; Schuler, B. Charge Interactions Can Dominate the Dimensions of Intrinsically Disordered Proteins. Proc. Natl. Acad. Sci. U. S. A. 2010, 107 (33), 14609–14614.

(75) Basu, S.; Martínez-Cristóbal, P.; Pesarrodona, M.; Frigolé-Vivas, M.; Lewis, M.; Szulc, E.; Bañuelos, C. A.; Sánchez-Zarzalejo, C.; Bielskutė, S.; Zhu, J.; Pombo-García, K.; Garcia-Cabau, C.; Batlle, C.; Mateos, B.; Biesaga, M.; Escobedo, A.; Bardia, L.; Verdaguer, X.; Ruffoni, A.; Mawji, N. R.; Wang, J.; Tam, T.; Brun-Heath, I.; Ventura, S.; Meierhofer, D.; García, J.; Robustelli, P.; Stracker, T. H.; Sadar, M. D.; Riera, A.; Hnisz, D.; Salvatella, X. Rational Optimization of a Transcription Factor Activation Domain Inhibitor. bioRxiv, 2022. https://doi.org/10.1101/2022.08.18.504385.

(76) Gomes, G.-N. W.; Krzeminski, M.; Namini, A.; Martin, E. W.; Mittag, T.; Head-Gordon, T.; Forman-Kay, J. D.; Gradinaru, C. C. Conformational Ensembles of an Intrinsically Disordered Protein Consistent with NMR, SAXS, and Single-Molecule FRET. J. Am. Chem. Soc. 2020, 142 (37), 15697–15710.

(77) Gibbs, E. B.; Lu, F.; Portz, B.; Fisher, M. J.; Medellin, B. P.; Laremore, T. N.; Zhang, Y. J.; Gilmour, D. S.; Showalter, S. A. Phosphorylation Induces Sequence-Specific Conformational Switches in the RNA Polymerase II C-Terminal Domain. Nat. Commun. 2017, 8 (1), 15233.

(78) Portz, B.; Lu, F.; Gibbs, E. B.; Mayfield, J. E.; Rachel Mehaffey, M.; Zhang, Y. J.; Brodbelt, J. S.; Showalter, S. A.; Gilmour, D. S. Structural Heterogeneity in the Intrinsically Disordered RNA Polymerase II C-Terminal Domain. Nat. Commun. 2017, 8, 15231.

(79) Wicky, B. I. M.; Shammas, S. L.; Clarke, J. Affinity of IDPs to Their Targets Is Modulated by Ion-Specific Changes in Kinetics and Residual Structure. Proc. Natl. Acad. Sci. U. S. A. 2017, 114 (37), 9882–9887.

(80) Moses, D.; Guadalupe, K.; Yu, F.; Flores, E.; Perez, A.; McAnelly, R.; Shamoon, N. M.; Cuevas-Zepeda, E.; Merg, A. D.; Martin, E. W.; Holehouse, A. S.; Sukenik, S. Structural Biases in Disordered Proteins Are Prevalent in the Cell. bioRxiv, 2022, 2021.11.24.469609. https://doi.org/10.1101/2021.11.24.469609.

(81) Naudi-Fabra, S.; Tengo, M.; Jensen, M. R.; Blackledge, M.; Milles, S. Quantitative Description of Intrinsically Disordered Proteins Using Single-Molecule FRET, NMR, and SAXS. J. Am. Chem. Soc. 2021, 143 (48), 20109–20121.

(82) Guseva, S.; Milles, S.; Jensen, M. R.; Salvi, N.; Kleman, J.-P.; Maurin, D.; Ruigrok, R. W. H.; Blackledge, M. Measles Virus Nucleo- and Phosphoproteins Form Liquid-like Phase-Separated Compartments That Promote Nucleocapsid Assembly. Sci Adv 2020, 6 (14), eaaz7095.

(83) Prestel, A.; Wichmann, N.; Martins, J. M.; Marabini, R.; Kassem, N.; Broendum, S. S.; Otterlei, M.; Nielsen, O.; Willemoës, M.; Ploug, M.; Boomsma, W.; Kragelund, B. B. The PCNA Interaction Motifs Revisited: Thinking Outside the PIP-Box. Cell. Mol. Life Sci. 2019. https://doi.org/10.1007/s00018-019-03150-0.

(84) Bugge, K.; Brakti, I.; Fernandes, C. B.; Dreier, J. E.; Lundsgaard, J. E.; Olsen, J. G.; Skriver, K.; Kragelund, B. B. Interactions by Disorder - A Matter of Context. Front Mol Biosci 2020, 7, 110.

(85) Cubuk, J.; Alston, J. J.; Incicco, J. J.; Singh, S.; Stuchell-Brereton, M. D.; Ward, M. D.; Zimmerman, M. I.; Vithani, N.; Griffith, D.; Wagoner, J. A.; Bowman, G. R.; Hall, K. B.; Soranno, A.; Holehouse, A. S. The SARS-CoV-2 Nucleocapsid Protein Is Dynamic, Disordered, and Phase Separates with RNA. Nat. Commun. 2021, 12 (1), 1936.

(86) Sankaranarayanan, M.; Emenecker, R. J.; Wilby, E. L.; Jahnel, M.; Trussina, I. R. E. A.; Wayland, M.; Alberti, S.; Holehouse, A. S.; Weil, T. T. Adaptable P Body Physical States Differentially Regulate Bicoid mRNA Storage during Early Drosophila Development. Dev. Cell 2021, 56 (20), 2886–2901.e6.

(87) Moses, D.; Guadalupe, K.; Yu, F.; Flores, E.; Perez, A.; McAnelly, R.; Shamoon, N. M.; Cuevas-Zepeda, E.; Merg, A.; Martin, E. W.; Holehouse, A. S.; Sukenik, S. Hidden Structure in Disordered Proteins Is Adaptive to Intracellular Changes. bioRxiv, 2021, 2021.11.24.469609. https://doi.org/10.1101/2021.11.24.469609.

(88) Husic, B. E.; Charron, N. E.; Lemm, D.; Wang, J.; Pérez, A.; Majewski, M.; Krämer, A.; Chen, Y.; Olsson, S.; de Fabritiis, G.; Noé, F.; Clementi, C. Coarse Graining Molecular Dynamics with Graph Neural Networks. arXiv [physics.comp-ph], 2020. http://arxiv.org/abs/2007.11412.

(89) Parton, D. L.; Grinaway, P. B.; Hanson, S. M.; Beauchamp, K. A.; Chodera, J. D. Ensembler: Enabling High-Throughput Molecular Simulations at the Superfamily Scale. PLoS Comput. Biol. 2016, 12 (6), e1004728.

(90) Singh, S.; Bowman, G. R. Quantifying Allosteric Communication via Both Concerted Structural Changes and Conformational Disorder with CARDS. J. Chem. Theory Comput. 2017, 13 (4), 1509–1517.

(91) Tubiana, T.; Carvaillo, J.-C.; Boulard, Y.; Bressanelli, S. TTClust: A Versatile Molecular Simulation Trajectory Clustering Program with Graphical Summaries. J. Chem. Inf. Model. 2018, 58 (11), 2178–2182.

(92) Lotz, S. D.; Dickson, A. Wepy: A Flexible Software Framework for Simulating Rare Events with Weighted Ensemble Resampling. ACS Omega 2020, 5 (49), 31608–31623.

(93) Richard, R. M.; Bertoni, C.; Boschen, J. S.; Keipert, K.; Pritchard, B.; Valeev, E. F.; Harrison, R. J.; de Jong, W. A.; Windus, T. L. Developing a Computational Chemistry Framework for the Exascale Era. Computing in Science Engineering 2019, 21 (2), 48–58.

(94) Meng, W.; Lyle, N.; Luan, B.; Raleigh, D. P.; Pappu, R. V. Experiments and Simulations Show How Long-Range Contacts Can Form in Expanded Unfolded Proteins with Negligible Secondary Structure. Proc. Natl. Acad. Sci. U. S. A. 2013, 110 (6), 2123–2128.

