## Supplementary Information for "SOURSOP: A Python package for the analysis of simulations of intrinsically disordered proteins"

### SUPPLEMENTARY METHODS

##### Excluded Volume (EV) simulations

As done previously [1,2], EV simulations were performed using the CAMPARI simulation engine and scaling all non-bonded interactions other than the repulsive component of the Lennard-Jones potential to zero. All EV simulation trajectories can be found in the trajectories link, found under the main GitHub repository for this manuscript at https://github.com/holehouse-lab/supportingdata/tree/master/2023/lalmansingh_2023.

### SUPPLEMENTARY TEXT

##### 1. The modern ecosystem of simulation analysis packages

The algorithmic power of trajectory analysis software dictates the extent to which biophysical insights can be gleaned from simulation trajectories. Broadly, such software can be categorized into four groups that are organized by the knowledge required for operational use as well as the degree of extensibility: frameworks, toolkits, plugins, and web servers (**Figure S1a**).

As subgroups draw functionalities from other groups or can be dependencies, a directed graph is shown in **Figure S1b** that further illustrates how the groups are interrelated. Here we provide an overview of each group ordered by user-friendliness and how SOURSOP fits into this ecosystem.

##### *1.1 Web servers*

In the biophysical community, web servers are services that abstract the functionality of a toolkit, plugin, or framework for use via a web interface such as a browser. Such services can perform specific calculations such as sequence alignments, trajectory analysis, etc. and report them to the user without the need for the installation of external software packages or tools.

Services that perform analysis of molecular trajectories typically only require the upload of a simulation trajectory and entry of select analysis options via form controls. Analysis is performed by the server using user-specified options and results are reported to the user via the browser. In cases where visualizations are output, these are presented as images for static plots while interactive visualizations can be presented via WebGL and JavaScript plugins such as NGLView and Three.js [3,4].

In this paradigm, the analysis functionality is performed via the use of frameworks, toolkits, plugins, or custom scripts utilizing mixtures of the three. Consequently, web servers typically adapt different programming languages and technology stacks to implement its functionality. Consequently, although web servers are easy to use and familiar, they can be more difficult to extend.

##### *1.2 Plugins*

In computing parlance, plugins are programs that extend the functionality of an existing piece of software. For trajectory analysis, plugins are programs or scripts which provide an additional set of analysis routines to a visualization toolkit such as VMD [5] or PyMol [6]. In some cases, plugins can significantly augment features of the host software, as in the case of Geo-Measures [7], which enables PyMol to analyze trajectories and perform ensemble analysis.

The major benefit of this approach is that the user can perform interactive analysis within a familiar interface without deep knowledge of the implementation of the host software or plugin. Typically, plugin-specific options are entered via input dialogs or through a text interface in the host software when required. To maximize interoperability, plugins are typically built in similar programming languages as the host software or are implemented in supported scripting languages.

Consequently, plugins can provide an easier path to the development of existing features for a given toolkit without the requirement of extensive knowledge of the host software's inner workings and implementation.

##### *1.3 Toolkits*

Within a computational biophysical context, toolkits are collections of standalone programs or scripts that can perform molecular simulations and related analyses. In some cases, these utilities can combine several routines into one interface, or are authored without the reliance on a given application framework. As such tools are typically launched via a command-line interface, analysis options can be passed directly as arguments to the specific program. This feature provides versatility that enables the rapid development of batch-processing scripts for analysis via the use of scripting programming languages.

A notable caveat about the programs within toolkits is that they can be difficult to extend, based on their implementation. Programs that are compiled can only be extended by modifying the underlying source code and recompiling, which can be a daunting and laborious process. However, toolkits derived from frameworks that use scripting programming languages such as Python have the greatest potential for extensibility. In such cases, new functionality can be added or implemented within executables by encapsulating key framework functions into a command-line interface.

##### *1.4 Frameworks*

Frameworks provide an application programming interface (API) in the chosen programming language of choice for the development of custom analysis functions, pipelines, and tools. Most frameworks are built with specific traits in mind, such as ease of access to specific libraries, ease of development based on a programming language's expressivity, or performance, to name but a few.

For example, frameworks developed in interpreted languages such as Python and R enable rapid development but can have lower runtime performance. However, the application of optimization libraries or packages that interface with compiled languages can alleviate such concerns. APIs provided in statically-typed programming languages such as C, C++, Fortran, and Java produce faster programs but require more time for development and are generally more verbose.

An important characteristic of frameworks is the level of detail they provide for the development of new functionality and analysis algorithms. This aspect is also intrinsically tied to the granularity of structural and representational details of a system's components, as they can enable or impede the kind of analyses that can be performed. For example, end-to-end distance calculations of proteins require access to the positions of the atoms within each residue and can be performed using the alpha Carbon or even the residue's center of mass. If individual residues are only implemented as a single [X, Y, Z] coordinate rather than collections of atoms, only the COM end-to-end distance can be calculated.

Consequently, several frameworks organize trajectory data into natural hierarchical objects such as atoms, amino acids, proteins, protein chains, and molecules. Some of these frameworks, such as MDTraj and MDAnalysis, adapt that architecture to provide powerful domain-specific languages for molecular selection, which can further enable even more rapid development of algorithms at different representational resolutions. Frameworks of interest that can be used in that context include MDTraj [8], MDAnalysis [9,10], LOOS [11], Jgromacs [12], OpenStructure [13], Pteros [14], CPPTraj [15], and Bio3D [16]. Thus, such frameworks serve as natural starting points for developing new frameworks and toolkits with specific use cases in mind.

Hence, the following frameworks descend from MD Analysis: PyTim [17], HiMach [18], Scoria [19], and Pteros [14]. For toolkits, the following packages employ MD Analysis for analysis routines or processing input files: HeroMDAnalysis [20], ProtoMD [21], PyContact [22], and PBxplore [23].

MDTraj is also represented in this context. For toolkits, the SSTMap [24], taurenmd [25], and TTClust [26] toolkits employ MDTraj for analysis and input processing. At the time of this writing, the frameworks Geo-Measures [7], MDEntropy [27], PyRETIS 2 [28], OpenPathSampling[29,30], and ProLint [31] employ MDTraj in their APIs. The Biotite package [32] is notable as its most recent version integrates both MDAnalysis and MDTraj.

In Supplementary **Table S1**, we survey a number of software packages that have been developed for the analysis of MD trajectories in different contexts. The majority of these packages are toolkits for quickly processing data and extracting relevant spatial properties as well as ensemble calculations that are important to molecular recognition and docking, molecular transport pathways, etc.

#### 2. Additional analyses provided in SOURSOP

SOURSOP allows for a wide range of analysis routines not described in this manuscript but that are outlined in the online documentation. In particular, SOURSOP natively facilitates multi-chain analysis, allowing interactions between and among distinct polypeptide chains to be analyzed.

As an example, interchain distance analysis is performed for GCN4, as taken from Robustelli et al. [33]. This analysis shows the inter-residue distances (in nm) between two GCN4 protein chains of the same length, although distance map calculations derived from chains of dissimilar lengths are also supported (**Fig. S1**). Distance maps can be computed using either a residue's α-carbon or the center of mass.

#### Supplemental Figures

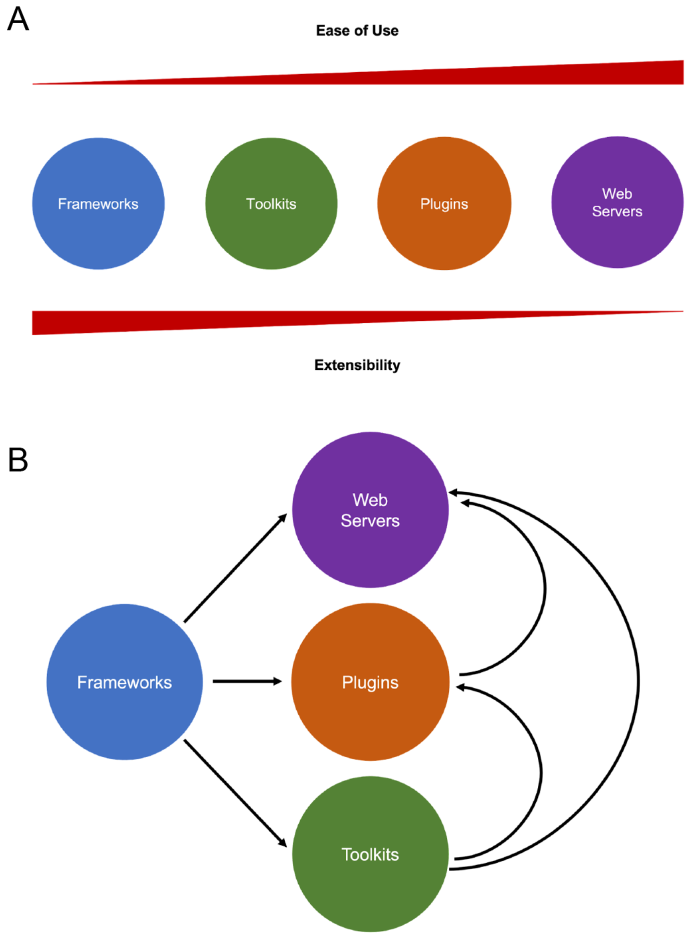


**Figure S1**: **Overview of trajectory analysis software categories and their interdependencies. (A)** An overview of the four trajectory analysis software categories that illustrate respective ease of use and extensibility. As software abstractions increase, utility and user-friendliness also increase. However, such abstractions decrease extensibility and interoperability with other software. **(B)** Dependencies and relationships among the four trajectory analysis software categories. Here, an arrow indicates a dependency relationship. Of these, Frameworks are the most versatile as they can be developed into any of the other categories.

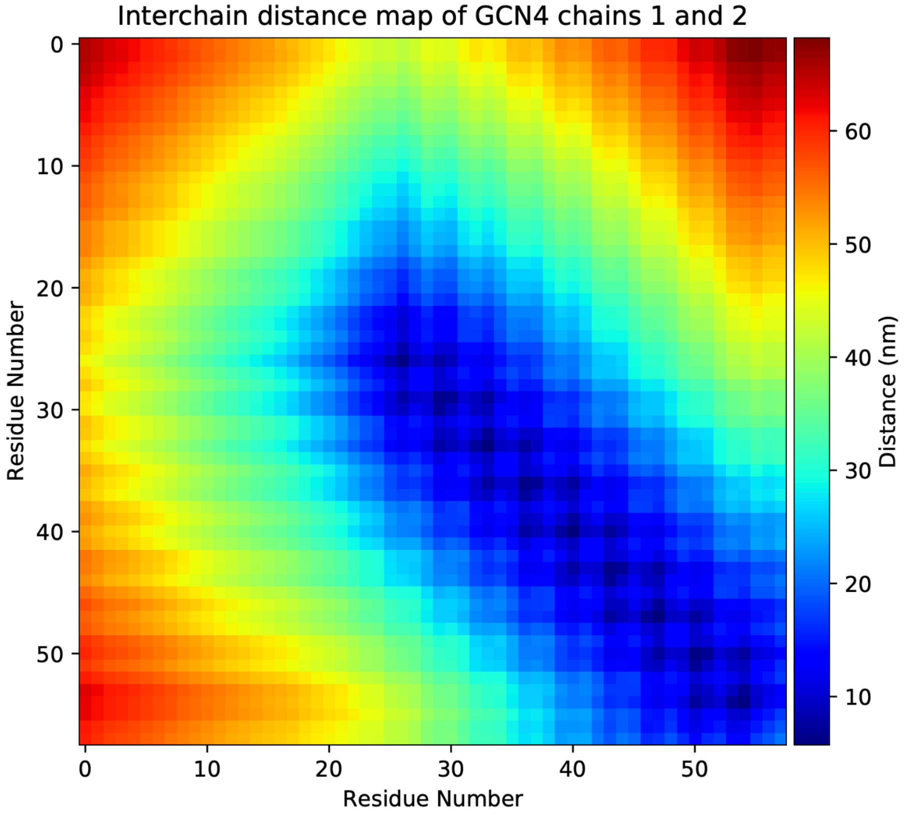


**Figure S2**: Intermolecular distance maps quantify interchain distances between two GCN4 protein chains in a dimeric configuration. This analysis shows that the majority of residues above index 20 on both chains are relatively close to one another. That suggests that those residues on both chains have formed and maintained stable intermolecular interactions through the extent of the simulation. Other chain residues appear to adopt a number of other conformations. We can further query the type of secondary structure formed by examining each chain's local collapse.

#### Supplemental Tables

| Software | Type | Software Use For Trajectory Analysis | Language | Trajectories | Year |
|---|---|---|---|---|---|
| nMoldyn [34] | Toolkit | Computation and decomposition of neutron scattering intensities | Fortran | MD | 1995 |
| HOLE [35] | Toolkit | Analysis of pore dimensions in ion channel structural models | Fortran | MD | 1996 |
| VMD [5] | Toolkit | Modeling, Visualization, & Structural Analysis | C | MD / MC | 1996 |
| MMTK [36] | Framework | Modeling & Structural Analysis | Python | MD | 2000 |
| PyMol [6] | Toolkit | Modeling, Visualization, & Structural Analysis | Python | MD / MC | 2002 |
| CAVER [37,38] | Toolkit | Analysis and clustering of transport pathways | C++ | MD | 2006 |
| MOLE [37] | Toolkit / Plugin / WebServer | Location and characterization of molecular channels, pores, and tunnels | Various | MD | 2007 |
| Wordom [39] | Toolkit | PCA, Entropy, Clustering, Conversion Tools | C | MD | 2007 |
| HiMach [18] | Framework | Distributed Trajectory Analysis | Python | MD | 2008 |
| Hollow [40] | Toolkit | Analysis of channel and interior surfaces in molecular structures | Python | MD | 2008 |
| MolAxis [41] | Toolkit / Webserver | Channel identification in macromolecules | Various | MD | 2008 |
| MSMBuilder [42] | Framework | Markov-State Model toolkit, with Spatial and Structural Analysis | Python / Scripts | MD | 2009 |
| PLUMED [43] | Plugin | Enhanced Sampling and Collective Variables | C | MD | 2009 |
| SMOG [44] | WebServer | CG structure-based modeling | Various | MD | 2010 |
| MAVENs [45] | Toolkit | ENMs for all-atom, united-atom, and CG trajectories | Matlab / Perl / C++ | MD | 2011 |
| POVME [46] | Toolkit | Pocket selection for ensemble docking | Python | MD | 2011 |
| ProDy [47] | Framework | Structural and Spatial analysis, with Visualization | Python | MD | 2011 |
| GROMOS++ [48] | Toolkit | Spatial, Structural, and information Analyses of GROMACS trajectories | C++ | MD | 2011 |
| TRAVIS [49] | Toolkit | Spatial, Structural, and Temporal Trajectory Analyses | C++ | MD / MC | 2011 |
| MDpocket [50] | Toolkit | Tracking small molecule binding sites and gas migration pathways | C | MD | 2011 |
| Eucb [51] | Toolkit | Identifies rare events by geometric criteria | C++ | MD | 2011 |
| MDAnalysis [9] | Framework | Spatial and Structural analysis | Python | MD | 2011 |
| GMCT [52] | Toolkit | Free energy calculations | C/C++ | MC | 2012 |
| MMPBSA.py [53] | Toolkit | Free energy calculations | Python | MD / MC | 2012 |
| Jgromacs [12] | Framework | Object-Oriented Representation and Analysis | Java | MD | 2012 |
| Pteros [54] | Framework | Geometry transformations, Structural alignment, Energy calculations | C++ | MD | 2012 |
| APL@Voro [55] | Toolkit | Analyze GROMACS lipid bilayer simulations, with Visualization | C++ | MD | 2013 |
| grcarma [56] | Toolkit | Spatial, Structural, and information Theoretic Analyses, | Perl / Tk | MD | 2013 |

| | | | | | |
|---|---|---|---|---|---|
| | | with Visualization | | | |
| Magic [57] | Framework | Structure based CG and analysis | Fortran / Python | MD / Inverse Monte Carlo | 2013 |
| OpenStructure [13] | Framework | Spatial, Structural, and information Theoretic Analyses, with Visualization | C++ & Python bindings | MD | 2013 |
| PTRAJ / CPPTRAJ / PYTRAJ [15] | Framework & Toolkit | Spatial, Structural, and information Theoretic Analyses | C++ | MD | 2013 |
| ReaDDy [58] | Framework | Analyze reaction diffusion dynamics in crowded environments | Java | MD | 2013 |
| Bio3D [16] | Framework | Spatial, Structural, Evolutionary Analysis, with Visualization | R | MD | 2014 |
| LOOS [11] | Framework | Spatial, Structural, and information Theoretic Analyses | C++ & Python bindings | MD | 2014 |
| ST-Analyzer [59] | WebServer | Spatial and Structural analyses for proteins and lipids | Python | MD | 2014 |
| Epock [60] | Toolkit & Plugin | Calculate pocket volumes, with visualization | C++ / Python / TCL | MD | 2015 |
| MDN [61] | WebServer | Characterize allosteric signaling pathways using ENMs | Various | MD | 2015 |
| MDTraj [8] | Framework | Spatial and Structural analysis | Python | MD / MC | 2015 |
| QwikMD [62] | Toolkit & Plugin | Spatial and Structural analyses, with Visualization | Various | MD | 2016 |
| ProtPOS [63] | Toolkit | Predict low-energy protein orientations on a surface of interest | Python | MD | 2016 |
| mDCC_tools [64] | Toolkit | Network Analysis, Pattern Recognition, with Visualization | Python Scripts and Gromacs Scripts | MD | 2016 |
| HTMD [65] | Framework | Protein Folding analysis and adaptive sampling ligand binding | Python | MD | 2016 |
| Parent [66] | Toolkit | Parallelized calculation of configurational entropy | C++ | MD | 2016 |
| pyPcazip [67] | Toolkit | Parallelized PCA analysis | Python | MD | 2016 |
| ProtoMD [21] | Framework & Toolkit | Spatial and Structural analyses, which can be used with CG-MD | Python | MD | 2016 |
| Scoria [19] | Framework | Simple molecular modeling with spatial analysis | Python | MD | 2017 |
| PBxplore [23] | Toolkit | Analyze the dynamics and deformability of protein structures | Python | MD | 2017 |
| MD-TASK [68] | Toolkit & Plugin | Performs perturbation response scanning, dynamic cross-correlation, and graph theory | Python | MD | 2017 |
| MDEntropy [27] | Framework | Information Theoretic analysis | Python | MD | 2017 |
| PyRETIS [69] | Toolkit | Identifies rare events by replica exchange | Python | MD | 2017 |
| Pytim [17] | Framework | Interfacial analysis and clustering, with Visualization in NGLView | Python | MD | 2018 |
| StreaMD [70] | Framework | Spatial, structural analysis, network transformations and analysis (CG supported) | R / C++ / Java | MD | 2018 |
| gRINN [71] | Toolkit | Residue Interaction Energies and Protein Energy Network analysis | Python | MD | 2018 |
| MODE-TASK [72] | Toolkit & Plugin | Large-Scale Protein motion analysis and visualization | C++ & Python bindings | MD | 2018 |
| RapidRMSD [73] | Framework & Toolkit | Rapid calculation of RMSDs for flexible molecules | C++ | MD | 2018 |
| SSTMap [24] | Toolkit | Thermodynamic, Structural, and Spatial Analysis of Water in MD trajectories | Python | MD | 2018 |

| | | | | | |
|---|---|---|---|---|---|
| TTClust [26] | Toolkit | Identify Dissimilar conformations through trajectory clustering | Python | MD | 2018 |
| PyContact [22] | Toolkit | Identify noncovalent interactions, with Visualization | Python | MD | 2018 |
| OpenPathSampling [30] | Framework | Path Sampling | Python | MD | 2018 |
| Biotite [32] | Framework | Spatial and Structural analysis | Python | MD | 2018 |
| SNP2SIM [74] | Toolkit | Predict binding to small molecule ligands | Python | MD | 2019 |
| NAPS [75] | WebServer | Network analysis based on inter-residue interaction energies | Mostly Python & PHP | MD | 2019 |
| ENSPARA [76] | Framework | Spatial and Structural analysis | Python | MD | 2019 |
| MERMAID [77] | WebServer | Prepare, run, and analyze CG MD models | PHP / Bash / JS | MD | 2019 |
| PCAViz [78] | Toolkit | PCA analysis and Visualization | Python / JS | MD | 2019 |
| Gromaps [79] | Toolkit | Computes and compares density maps of MD simulation trajectories | Various | MD | 2019 |
| AViS [80] | Toolkit | Dataflow programming implementation for analyzing MD trajectories, with visualization | OpenGL / C++ | MD | 2020 |
| Geo-Measures [7] | Plugin | PyMol plugin for structure ensemble analysis | Python | MD | 2020 |
| Protlego [81] | Framework | Design and structural analysis of chimeric proteins | Python | MD | 2020 |
| PyVisA [82] | Toolkit | Visualization and analysis of path sampling trajectories | Python | MD | 2020 |
| sDMD [83] | Toolkit | Structural and Spatial for Discontinuous MD | C/C++ | MD | 2020 |
| taurenmd [25] | Toolkit | Spatial and Structural analysis | Python | MD | 2020 |
| quickSOM [84] | Toolkit | Cluster the conformational ensembles of MD simulations | Python | MD | 2021 |
| HeroMDAnalysis [20] | Toolkit | Toolset for the rapid analysis and visualization of GROMACS trajectories | Bash & Zenity | MD | 2021 |
| JEDi [85] | Toolkit | Parallelized structural and spatial analysis | Java | MD | 2021 |
| ProLint [31] | Framework & Web server | Analysis and visualization for lipid-protein interactions | Python | MD | 2021 |

**Supplemental Table 1*:* A survey of trajectory analysis software packages with significant functions implemented and updated in the past 20 years.** Shown is a list of major software packages that perform trajectory analysis ordered by the year when it was first mentioned in the literature. Although Toolkits and Frameworks are represented the most, there are several plugins for visualization and analysis programs that could have also been included. The majority of these were excluded as they did not provide a significant function set that was not encapsulated in another package. In some cases, the software was significantly updated, as in the case of CAVER [38,86,87], nMoldyn [34,88,89], PLUMED [43,90], Pteros [14,54], SMOG [44,91], POVME [46,92,93], MDAnalysis [9,10], HOLE [35,94], MOLE [37,95], and ReaDDy [58,96]. In other instances, some software packages were incrementally updated, but their updates were not officially published via the literature.

#### SUPPLEMENTAL REFERENCES


(1) Holehouse, A. S.; Garai, K.; Lyle, N.; Vitalis, A.; Pappu, R. V. Quantitative Assessments of the Distinct Contributions of Polypeptide Backbone Amides versus Side Chain Groups to Chain Expansion via Chemical Denaturation. *J. Am. Chem. Soc.* **2015**, *137* (8), 2984–2995.

(2) Mao, A. H.; Crick, S. L.; Vitalis, A.; Chicoine, C. L.; Pappu, R. V. Net Charge per Residue Modulates Conformational Ensembles of Intrinsically Disordered Proteins. *Proc. Natl. Acad. Sci. U. S. A.* **2010**, *107* (18), 8183–8188.

(3) Nguyen, H.; Case, D. A.; Rose, A. S. NGLview–interactive Molecular Graphics for Jupyter Notebooks. *Bioinformatics* **2017**, *34* (7), 1241–1242.

(4) Danchilla, B. Three.js Framework. In *Beginning WebGL for HTML5*; Danchilla, B., Ed.; Apress: Berkeley, CA, 2012; pp 173–203.

(5) Humphrey, W.; Dalke, A.; Schulten, K. VMD: Visual Molecular Dynamics. *J. Mol. Graph.* **1996**, *14* (1), 33–38, 27–28.

(6) DeLano, W. L.; Others. Pymol: An Open-Source Molecular Graphics Tool. *CCP4 Newsletter on protein crystallography* **2002**, *40* (1), 82–92.

(7) Kagami, L. P.; das Neves, G. M.; Timmers, L. F. S. M.; Caceres, R. A.; Eifler-Lima, V. L. Geo-Measures: A PyMOL Plugin for Protein Structure Ensembles Analysis. *Comput. Biol. Chem.* **2020**, *87*, 107322.

(8) McGibbon, R. T.; Beauchamp, K. A.; Harrigan, M. P.; Klein, C.; Swails, J. M.; Hernández, C. X.; Schwantes, C. R.; Wang, L.-P.; Lane, T. J.; Pande, V. S. MDTraj: A Modern, Open Library for the Analysis of Molecular Dynamics Trajectories. *Biophys. J.* **2015**, *109* (8), 1528–1532.

(9) Michaud-Agrawal, N.; Denning, E. J.; Woolf, T. B.; Beckstein, O. MDAnalysis: A Toolkit for the Analysis of Molecular Dynamics Simulations. *J. Comput. Chem.* **2011**, *32* (10), 2319–2327.

(10) Gowers, R. J.; Linke, M.; Barnoud, J.; Reddy, T. J. E.; Melo, M. N. MDAnalysis: A Python Package for the Rapid Analysis of Molecular Dynamics Simulations. **2019**.

(11) Romo, T. D.; Leioatts, N.; Grossfield, A. Lightweight Object Oriented Structure Analysis: Tools for Building Tools to Analyze Molecular Dynamics Simulations. *J. Comput. Chem.* **2014**, *35* (32), 2305–2318.

(12) Münz, M.; Biggin, P. C. JGromacs: A Java Package for Analyzing Protein Simulations. *J. Chem. Inf. Model.* **2012**, *52* (1), 255–259.

(13) Biasini, M.; Schmidt, T.; Bienert, S.; Mariani, V.; Studer, G.; Haas, J.; Johner, N.; Schenk, A. D.; Philippsen, A.; Schwede, T. OpenStructure: An Integrated Software Framework for Computational Structural Biology. *Acta Crystallogr. D Biol. Crystallogr.* **2013**, *69* (Pt 5), 701–709.

(14) Yesylevskyy, S. O. Pteros 2.0: Evolution of the Fast Parallel Molecular Analysis Library for C++ and Python. *J. Comput. Chem.* **2015**, *36* (19), 1480–1488.

(15) Roe, D. R.; Cheatham, T. E., 3rd. PTRAJ and CPPTRAJ: Software for Processing and Analysis of Molecular Dynamics Trajectory Data. *J. Chem. Theory Comput.* **2013**, *9* (7), 3084–3095.

(16) Skjærven, L.; Yao, X.-Q.; Scarabelli, G.; Grant, B. J. Integrating Protein Structural Dynamics and Evolutionary Analysis with Bio3D. *BMC Bioinformatics* **2014**, *15*, 399.

(17) Sega, M.; Hantal, G.; Fábián, B.; Jedlovszky, P. Pytim: A Python Package for the Interfacial Analysis of Molecular Simulations. *J. Comput. Chem.* **2018**, *39* (25), 2118–2125.

(18) Tu, T.; Rendleman, C. A.; Borhani, D. W.; Dror, R. O.; Gullingsrud, J.; Jensen, M. O.; Klepeis, J. L.; Maragakis, P.; Miller, P.; Stafford, K. A.; Shaw, D. E. A Scalable Parallel Framework for Analyzing Terascale Molecular Dynamics Simulation Trajectories. In *SC '08: Proceedings of the 2008 ACM/IEEE Conference on Supercomputing*; ieeexplore.ieee.org,

2008; pp 1–12.
(19) Ropp, P.; Friedman, A.; Durrant, J. D. Scoria: A Python Module for Manipulating 3D Molecular Data. *J. Cheminform.* **2017**, *9* (1), 52.
(20) Rawat, R.; Kant, K.; Kumar, A.; Bhati, K.; Verma, S. M. HeroMDAnalysis: An Automagical Tool for GROMACS-Based Molecular Dynamics Simulation Analysis. *Future Med. Chem.* **2021**, *13* (5), 447–456.
(21) Somogyi, E.; Mansour, A. A.; Ortoleva, P. J. ProtoMD: A Prototyping Toolkit for Multiscale Molecular Dynamics. *Comput. Phys. Commun.* **2016**, *202*, 337–350.
(22) Scheurer, M.; Rodenkirch, P.; Siggel, M.; Bernardi, R. C.; Schulten, K.; Tajkhorshid, E.; Rudack, T. PyContact: Rapid, Customizable, and Visual Analysis of Noncovalent Interactions in MD Simulations. *Biophys. J.* **2018**, *114* (3), 577–583.
(23) Barnoud, J.; Santuz, H.; Craveur, P.; Joseph, A. P.; Jallu, V.; de Brevern, A. G.; Poulain, P. PBxplore: A Tool to Analyze Local Protein Structure and Deformability with Protein Blocks. *PeerJ* **2017**, *5*, e4013.
(24) Haider, K.; Cruz, A.; Ramsey, S.; Gilson, M. K.; Kurtzman, T. Solvation Structure and Thermodynamic Mapping (SSTMap): An Open-Source, Flexible Package for the Analysis of Water in Molecular Dynamics Trajectories. *J. Chem. Theory Comput.* **2018**, *14* (1), 418–425.
(25) Teixeira, J. Taurenmd: A Command-Line Interface for Analysis of Molecular Dynamics Simulations. *J. Open Source Softw.* **2020**, *5* (50), 2175.
(26) Tubiana, T.; Carvaillo, J.-C.; Boulard, Y.; Bressanelli, S. TTClust: A Versatile Molecular Simulation Trajectory Clustering Program with Graphical Summaries. *J. Chem. Inf. Model.* **2018**, *58* (11), 2178–2182.
(27) X. Hernández, C.; S. Pande, V. MDEntropy: Information-Theoretic Analyses for Molecular Dynamics. *J. Open Source Softw.* **2017**, *2* (19), 427.
(28) Riccardi, E.; Lervik, A.; Roet, S.; Aarøen, O.; van Erp, T. S. PyRETIS 2: An Improbability Drive for Rare Events. *J. Comput. Chem.* **2020**, *41* (4), 370–377.
(29) Swenson, D. W. H.; Prinz, J.-H.; Noe, F.; Chodera, J. D.; Bolhuis, P. G. OpenPathSampling: A Python Framework for Path Sampling Simulations. 2. Building and Customizing Path Ensembles and Sample Schemes. *J. Chem. Theory Comput.* **2019**, *15* (2), 837–856.
(30) Swenson, D. W. H.; Prinz, J.-H.; Noe, F.; Chodera, J. D.; Bolhuis, P. G. OpenPathSampling: A Python Framework for Path Sampling Simulations. 1. Basics. *J. Chem. Theory Comput.* **2019**, *15* (2), 813–836.
(31) Sejdiu, B. I.; Tieleman, D. P. ProLint: A Web-Based Framework for the Automated Data Analysis and Visualization of Lipid–protein Interactions. *Nucleic Acids Res.* **2021**, *49* (W1), W544–W550.
(32) Kunzmann, P.; Hamacher, K. Biotite: A Unifying Open Source Computational Biology Framework in Python. *BMC Bioinformatics* **2018**, *19* (1), 346.
(33) Robustelli, P.; Piana, S.; Shaw, D. E. Developing a Molecular Dynamics Force Field for Both Folded and Disordered Protein States. *Proc. Natl. Acad. Sci. U. S. A.* **2018**, *115* (21), E4758–E4766.
(34) nMOLDYN: A Program Package for a Neutron Scattering Oriented Analysis of Molecular Dynamics Simulations. *Comput. Phys. Commun.* **1995**, *91* (1-3), 191–214.
(35) HOLE: A Program for the Analysis of the Pore Dimensions of Ion Channel Structural Models. *J. Mol. Graph.* **1996**, *14* (6), 354–360.
(36) Hinsen, K. The Molecular Modeling Toolkit: A New Approach to Molecular Simulations. *J. Comput. Chem.* **2000**, *21* (2), 79–85.
(37) Petrek, M.; Kosinová, P.; Koca, J.; Otyepka, M. MOLE: A Voronoi Diagram-Based Explorer of Molecular Channels, Pores, and Tunnels. *Structure* **2007**, *15* (11), 1357–1363.
(38) Petrek, M.; Otyepka, M.; Banás, P.; Kosinová, P.; Koca, J.; Damborský, J. CAVER: A New Tool to Explore Routes from Protein Clefts, Pockets and Cavities. *BMC Bioinformatics*

**2006**, *7*, 316.
(39) Seeber, M.; Cecchini, M.; Rao, F.; Settanni, G.; Caflisch, A. Wordom: A Program for Efficient Analysis of Molecular Dynamics Simulations. *Bioinformatics* **2007**, *23* (19), 2625–2627.
(40) Ho, B. K.; Gruswitz, F. HOLLOW: Generating Accurate Representations of Channel and Interior Surfaces in Molecular Structures. *BMC Struct. Biol.* **2008**, *8* (1), 1–6.
(41) Yaffe, E.; Fishelovitch, D.; Wolfson, H. J.; Halperin, D.; Nussinov, R. MolAxis: Efficient and Accurate Identification of Channels in Macromolecules. *Proteins* **2008**, *73* (1), 72–86.
(42) Bowman, G. R.; Huang, X.; Pande, V. S. Using Generalized Ensemble Simulations and Markov State Models to Identify Conformational States. *Methods* **2009**, *49* (2), 197–201.
(43) PLUMED: A Portable Plugin for Free-Energy Calculations with Molecular Dynamics. *Comput. Phys. Commun.* **2009**, *180* (10), 1961–1972.
(44) Noel, J. K.; Whitford, P. C.; Sanbonmatsu, K. Y.; Onuchic, J. N. SMOG@ctbp: Simplified Deployment of Structure-Based Models in GROMACS. *Nucleic Acids Res.* **2010**, *38* (Web Server issue), W657–W661.
(45) Zimmermann, M. T.; Kloczkowski, A.; Jernigan, R. L. MAVENs: Motion Analysis and Visualization of Elastic Networks and Structural Ensembles. *BMC Bioinformatics* **2011**, *12*, 264.
(46) Durrant, J. D.; de Oliveira, C. A. F.; McCammon, J. A. POVME: An Algorithm for Measuring Binding-Pocket Volumes. *J. Mol. Graph. Model.* **2011**, *29* (5), 773–776.
(47) Bakan, A.; Meireles, L. M.; Bahar, I. ProDy: Protein Dynamics Inferred from Theory and Experiments. *Bioinformatics* **2011**, *27* (11), 1575–1577.
(48) Eichenberger, A. P.; Allison, J. R.; Dolenc, J.; Geerke, D. P.; Horta, B. A. C.; Meier, K.; Oostenbrink, C.; Schmid, N.; Steiner, D.; Wang, D.; van Gunsteren, W. F. GROMOS++ Software for the Analysis of Biomolecular Simulation Trajectories. *J. Chem. Theory Comput.* **2011**, *7* (10), 3379–3390.
(49) Brehm, M.; Kirchner, B. TRAVIS - a Free Analyzer and Visualizer for Monte Carlo and Molecular Dynamics Trajectories. *J. Chem. Inf. Model.* **2011**, *51* (8), 2007–2023.
(50) Schmidtke, P.; Bidon-Chanal, A.; Luque, F. J.; Barril, X. MDpocket: Open-Source Cavity Detection and Characterization on Molecular Dynamics Trajectories. *Bioinformatics* **2011**, *27* (23), 3276–3285.
(51) Tsoulos, I. G.; Stavrakoudis, A. Eucb: A C++ Program for Molecular Dynamics Trajectory Analysis. *Comput. Phys. Commun.* **2011**, *182* (3), 834–841.
(52) Ullmann, R. T.; Ullmann, G. M. GMCT : A Monte Carlo Simulation Package for Macromolecular Receptors. *J. Comput. Chem.* **2012**, *33* (8), 887–900.
(53) Miller, B. R., 3rd; McGee, T. D., Jr; Swails, J. M.; Homeyer, N.; Gohlke, H.; Roitberg, A. E. MMPBSA.py: An Efficient Program for End-State Free Energy Calculations. *J. Chem. Theory Comput.* **2012**, *8* (9), 3314–3321.
(54) Yesylevskyy, S. O. Pteros: Fast and Easy to Use Open-Source C++ Library for Molecular Analysis. *J. Comput. Chem.* **2012**, *33* (19), 1632–1636.
(55) Lukat, G.; Krüger, J.; Sommer, B. APL@Voro: A Voronoi-Based Membrane Analysis Tool for GROMACS Trajectories. *J. Chem. Inf. Model.* **2013**, *53* (11), 2908–2925.
(56) Koukos, P. I.; Glykos, N. M. Grcarma: A Fully Automated Task-Oriented Interface for the Analysis of Molecular Dynamics Trajectories. *J. Comput. Chem.* **2013**, *34* (26), 2310–2312.
(57) Mirzoev, A.; Lyubartsev, A. P. MagiC: Software Package for Multiscale Modeling. *J. Chem. Theory Comput.* **2013**, *9* (3), 1512–1520.
(58) Schöneberg, J.; Noé, F. ReaDDy - A Software for Particle-Based Reaction-Diffusion Dynamics in Crowded Cellular Environments. *PLoS One* **2013**, *8* (9), e74261.
(59) Jeong, J. C.; Jo, S.; Wu, E. L.; Qi, Y.; Monje-Galvan, V.; Yeom, M. S.; Gorenstein, L.; Chen, F.; Klauda, J. B.; Im, W. ST-Analyzer: A Web-Based User Interface for Simulation Trajectory Analysis. *J. Comput. Chem.* **2014**, *35* (12), 957–963.

(60) Laurent, B.; Chavent, M.; Cragnolini, T.; Dahl, A. C. E.; Pasquali, S.; Derreumaux, P.; Sansom, M. S. P.; Baaden, M. Epock: Rapid Analysis of Protein Pocket Dynamics. *Bioinformatics* **2015**, *31* (9), 1478–1480.
(61) Ribeiro, A. A. S. T.; Ortiz, V. MDN: A Web Portal for Network Analysis of Molecular Dynamics Simulations. *Biophys. J.* **2015**, *109* (6), 1110–1116.
(62) Ribeiro, J. V.; Bernardi, R. C.; Rudack, T.; Stone, J. E.; Phillips, J. C.; Freddolino, P. L.; Schulten, K. QwikMD — Integrative Molecular Dynamics Toolkit for Novices and Experts. *Sci. Rep.* **2016**, *6* (1), 1–14.
(63) Ngai, J. C. F.; Mak, P.-I.; Siu, S. W. I. ProtPOS: A Python Package for the Prediction of Protein Preferred Orientation on a Surface. *Bioinformatics* **2016**, *32* (16), 2537–2538.
(64) Kasahara, K.; Mohan, N.; Fukuda, I.; Nakamura, H. mDCC_tools: Characterizing Multi-Modal Atomic Motions in Molecular Dynamics Trajectories. *Bioinformatics* **2016**, *32* (16), 2531–2533.
(65) Doerr, S.; Harvey, M. J.; Noé, F.; De Fabritiis, G. HTMD: High-Throughput Molecular Dynamics for Molecular Discovery. *J. Chem. Theory Comput.* **2016**, *12* (4), 1845–1852.
(66) Fleck, M.; Polyansky, A. A.; Zagrovic, B. PARENT: A Parallel Software Suite for the Calculation of Configurational Entropy in Biomolecular Systems. *J. Chem. Theory Comput.* **2016**, *12* (4), 2055–2065.
(67) Shkurti, A.; Goni, R.; Andrio, P.; Breitmoser, E.; Bethune, I.; Orozco, M.; Laughton, C. A. pyPcazip: A PCA-Based Toolkit for Compression and Analysis of Molecular Simulation Data. *SoftwareX* **2016**, *5*, 44–50.
(68) Brown, D. K.; Penkler, D. L.; Sheik Amamuddy, O.; Ross, C.; Atilgan, A. R.; Atilgan, C.; Tastan Bishop, Ö. MD-TASK: A Software Suite for Analyzing Molecular Dynamics Trajectories. *Bioinformatics* **2017**, *33* (17), 2768–2771.
(69) Lervik, A.; Riccardi, E.; van Erp, T. S. PyRETIS: A Well-Done, Medium-Sized Python Library for Rare Events. *J. Comput. Chem.* **2017**, *38* (28), 2439–2451.
(70) Dombrowsky, M. J.; Jager, S.; Schiller, B.; Mayer, B. E.; Stammler, S.; Hamacher, K. StreaMD: Advanced Analysis of Molecular Dynamics Using R. *J. Comput. Chem.* **2018**, *39* (21), 1666–1674.
(71) Serçinoglu, O.; Ozbek, P. gRINN: A Tool for Calculation of Residue Interaction Energies and Protein Energy Network Analysis of Molecular Dynamics Simulations. *Nucleic Acids Res.* **2018**, *46* (W1), W554–W562.
(72) Ross, C.; Nizami, B.; Glenister, M.; Sheik Amamuddy, O.; Atilgan, A. R.; Atilgan, C.; Tastan Bishop, Ö. MODE-TASK: Large-Scale Protein Motion Tools. *Bioinformatics* **2018**, *34* (21), 3759–3763.
(73) Neveu, E.; Popov, P.; Hoffmann, A.; Migliosi, A.; Besseron, X.; Danoy, G.; Bouvry, P.; Grudinin, S. RapidRMSD: Rapid Determination of RMSDs Corresponding to Motions of Flexible Molecules. *Bioinformatics* **2018**, *34* (16), 2757–2765.
(74) McCoy, M. D.; Shivakumar, V.; Nimmagadda, S.; Jafri, M. S.; Madhavan, S. SNP2SIM: A Modular Workflow for Standardizing Molecular Simulation and Functional Analysis of Protein Variants. *BMC Bioinformatics* **2019**, *20* (1), 171.
(75) Chakrabarty, B.; Naganathan, V.; Garg, K.; Agarwal, Y.; Parekh, N. NAPS Update: Network Analysis of Molecular Dynamics Data and Protein-Nucleic Acid Complexes. *Nucleic Acids Res.* **2019**, *47* (W1), W462–W470.
(76) Porter, J. R.; Zimmerman, M. I.; Bowman, G. R. Enspara: Modeling Molecular Ensembles with Scalable Data Structures and Parallel Computing. *J. Chem. Phys.* **2019**, *150* (4), 044108.
(77) Damre, M.; Marchetto, A.; Giorgetti, A. MERMAID: Dedicated Web Server to Prepare and Run Coarse-Grained Membrane Protein Dynamics. *Nucleic Acids Res.* **2019**, *47* (W1), W456–W461.
(78) Pacheco, S.; Kaminsky, J. C.; Kochnev, I. K.; Durrant, J. D. PCAViz: An Open-Source

Python/JavaScript Toolkit for Visualizing Molecular Dynamics Simulations in the Web Browser. *J. Chem. Inf. Model.* **2019**, *59* (10), 4087–4092.
(79) Briones, R.; Blau, C.; Kutzner, C.; de Groot, B. L.; Aponte-Santamaría, C. Gromaps: A Gromacs-Based Toolset to Analyse Density Maps Derived from Molecular Dynamics Simulations. *Biophys. J.* **2019**, *116* (3), 142a – 143a.
(80) Pua, K.; Yuhara, D.; Ayuba, S.; Yasuoka, K. Dataflow Programming for the Analysis of Molecular Dynamics with AViS, an Analysis and Visualization Software Application. *PLoS One* **2020**, *15* (4), e0231714.
(81) Ferruz, N.; Noske, J.; Höcker, B. Protlego: A Python Package for the Analysis and Design of Chimeric Proteins. *Bioinformatics* **2021**. https://doi.org/10.1093/bioinformatics/btab253.
(82) Aarøen, O.; Kiaer, H.; Riccardi, E. PyVisA: Visualization and Analysis of Path Sampling Trajectories. *J. Comput. Chem.* **2021**, *42* (6), 435–446.
(83) Zheng, S.; Javidpour, L.; Sahimi, M.; Shing, K. S.; Nakano, A. sDMD: An Open Source Program for Discontinuous Molecular Dynamics Simulation of Protein Folding and Aggregation. *Comput. Phys. Commun.* **2020**, *247*, 106873.
(84) Mallet, V.; Nilges, M.; Bouvier, G. Quicksom: Self-Organizing Maps on GPUs for Clustering of Molecular Dynamics Trajectories. *Bioinformatics* **2021**, *37* (14), 2064–2065.
(85) David, C. C.; Avery, C. S.; Jacobs, D. J. JEDi: Java Essential Dynamics Inspector - a Molecular Trajectory Analysis Toolkit. *BMC Bioinformatics* **2021**, *22* (1), 226.
(86) Medek, P.; Beneš, P.; Kozlíková, B.; Chovancová, E.; Pavelka, A.; Szabó, T.; Zamborský, M.; Andres, F.; Klvaňa, M.; Brezovský, J.; Sochor, J.; Damborský, J. CAVER 2.0. **2008**.
(87) Chovancova, E.; Pavelka, A.; Benes, P.; Strnad, O.; Brezovsky, J.; Kozlikova, B.; Gora, A.; Sustr, V.; Klvana, M.; Medek, P.; Biedermannova, L.; Sochor, J.; Damborsky, J. CAVER 3.0: A Tool for the Analysis of Transport Pathways in Dynamic Protein Structures. *PLoS Comput. Biol.* **2012**, *8* (10), e1002708.
(88) Róg, T.; Murzyn, K.; Hinsen, K.; Kneller, G. R. nMoldyn: A Program Package for a Neutron Scattering Oriented Analysis of Molecular Dynamics Simulations. *J. Comput. Chem.* **2003**, *24* (5), 657–667.
(89) Hinsen, K.; Pellegrini, E.; Stachura, S.; Kneller, G. R. nMoldyn 3: Using Task Farming for a Parallel Spectroscopy-Oriented Analysis of Molecular Dynamics Simulations. *J. Comput. Chem.* **2012**, *33* (25), 2043–2048.
(90) Tribello, G. A.; Bonomi, M.; Branduardi, D.; Camilloni, C.; Bussi, G. PLUMED 2: New Feathers for an Old Bird. *Comput. Phys. Commun.* **2014**, *185* (2), 604–613.
(91) Noel, J. K.; Levi, M.; Raghunathan, M.; Lammert, H.; Hayes, R. L.; Onuchic, J. N.; Whitford, P. C. SMOG 2: A Versatile Software Package for Generating Structure-Based Models. *PLoS Comput. Biol.* **2016**, *12* (3), e1004794.
(92) Durrant, J. D.; Votapka, L.; Sørensen, J.; Amaro, R. E. POVME 2.0: An Enhanced Tool for Determining Pocket Shape and Volume Characteristics. *J. Chem. Theory Comput.* **2014**, *10* (11), 5047–5056.
(93) Wagner, J. R.; Sørensen, J.; Hensley, N.; Wong, C.; Zhu, C.; Perison, T.; Amaro, R. E. POVME 3.0: Software for Mapping Binding Pocket Flexibility. *J. Chem. Theory Comput.* **2017**, *13* (9), 4584–4592.
(94) Smart, O. *HOLE program homepage*. http://www.holeprogram.org/ (accessed 2022-11-14).
(95) Sehnal, D.; Svobodová Vařeková, R.; Berka, K.; Pravda, L.; Navrátilová, V.; Banáš, P.; Ionescu, C.-M.; Otyepka, M.; Koča, J. MOLE 2.0: Advanced Approach for Analysis of Biomacromolecular Channels. *J. Cheminform.* **2013**, *5* (1), 1–13.
(96) Hoffmann, M.; Fröhner, C.; Noé, F. ReaDDy 2: Fast and Flexible Software Framework for Interacting-Particle Reaction Dynamics. *PLoS Comput. Biol.* **2019**, *15* (2), e1006830.
